# Order matters: Autocorrelation of temperature dictates extinction risk in populations with nonlinear thermal performance

**DOI:** 10.1101/2024.12.19.629491

**Authors:** Alison J. Robey, David A. Vasseur

**Affiliations:** Department of Ecology & Evolutionary Biology, Yale University, P.O. Box 208106, New Haven, CT 06520-8106

## Abstract

Forecasting the risks caused by climate change often relies upon combining species’ thermal performance curves with expected statistical distributions of experienced temperatures, without consideration for the order in which those temperatures occur. Such averaging approaches may obscure the disproportionate impacts that extreme events like heatwaves have on fitness and survival. In this study, we instead incorporate thermal performance curves with population dynamical modeling to elucidate the relationship between the sequence of temperature events – driven by temporal autocorrelation – and extinction risk. We show that the permutation of temperatures determines the extent of risk; as thermal regimes grow warmer, more variable, and more autocorrelated, the risk of extinction grows non-linearly and is driven by interactions among our three treatment variables. Given that the mean, variance, and autocorrelation of temperatures are changing in nuanced ways across the globe, understanding these interactions is paramount for forecasting risk. Using empirical data from a benchmarked set of thermal performance curves, we demonstrate how extinction risk is impacted by the change in mean, variance, and autocorrelation, while controlling for seasonal and diurnal cycling. Our results and modeling approach offer new tools for testing the robustness of thermal performance curves and emphasize the importance of looking beyond temporally-blind metrics, like mean population size or average thermal distributions, for forecasting impending extinction risks.

## 1 Introduction

A species’ sensitivity to changing thermal environments depends both on how temperatures are changing and on its capacity to respond to those changes (Deutsch et al., 2008; Huey et al., 2012; Buckley et al., 2022). Different organisms, populations, and species have vastly different thermal sensitivities, dependent on the environments they have experienced, grown in, and evolved under (Kingsolver et al., 2013; Munday et al., 2013; Sinclair et al., 2016). The life or death of an individual – and the persistence or extinction of a broader population – is thus intrinsically dependent on its thermal history. Consequently, estimating thermal risk without accounting for the explicit ordering in which temperatures occur may not accurately capture the effects of increasing thermal variability and warming.

An increasingly popular method of forecasting thermal risk predicts long-term average performance or fitness by integrating a thermal performance curve (TPC) over a projected frequency distribution of temperatures (Huey and Stevenson, 1979; Deutsch et al., 2008; Tewksbury et al., 2008; Vasseur et al., 2014; Pinsky et al., 2019; Duffy et al., 2022). Although useful, this approach is based upon a mathematical averaging that does not account for the outsized impact catastrophic events like heatwaves have on individuals or populations. For example, a temporal sequence of highly stressful temperatures, like a series of exceptionally hot days during a heatwave, could cause death or extinction despite generating only a small reduction in the calculated long-term average fitness. While many improvements have been made to this method, such as incorporating high-resolution thermal data and adjusting TPCs to more accurately depict the severe consequences of extreme temperatures (e.g., Kingsolver et al., 2013; Buckley et al., 2022), its temporally-blind nature ultimately limits its utility. Improving our TPC-based risk forecasting requires us to move beyond such assumptions.

At the population level, risk is often attributed to population dynamic phenomena such as decline or extinction (Lande, 1993; Thomas et al., 2004; Urban, 2015). Because population dynamics are inherently sensitive to the temporal sequencing or ‘autocorrelation’ of environmental changes, merging TPCs into population dynamic models may provide a stronger basis for measuring risk than classical averaging approaches. Ecological theory has long recognized the potential for environmental temporal autocorrelation to increase extinction risk by clumping harmful events together (Lawton, 1988; Vasseur and Yodzis, 2004; Schwager et al., 2006). Physiologists have similarly found that temporal autocorrelation increases organism sensitivity to those stressful conditions (Rezende et al., 2014; Sinclair et al., 2016; Jørgensen et al., 2019; Rezende et al., 2020). In much of the world, projected intensification of heatwave regimes exacerbate such circumstances; using time-frequency decomposition of CMIP6 models, Duffy et al. (2022) found increasing temporal autocorrelation at 60% of aquatic and 80% of terrestrial locations (excluding Antarctica). The rising global incidence of stressful temperatures, where fitness and performance precipitously decline (Jørgensen et al., 2022), heightens the potential for warming thermal regimes to increase extinction risks in a manner dependent on the order in which those temperatures occur. Because current risk estimates broadly exclude explicit thermal regimes and their corresponding population dynamics, they are likely underestimating extinction and extirpation risks of species experiencing stressful, variable, and highly autocorrelated thermal regimes.

In this paper, we test the importance of incorporating population dynamics into forecasting models. While Duffy et al. (2022) offered an initial assessment of how population dynamics compare between historic and forecasted thermal regimes, they do not explicitly examine the effects of varying temporal autocorrelation. We advance this work by testing how population size, extinction risk, and subsequent risk metrics respond to variation in temporal autocorrelation across statistically identical thermal regimes (3.1); assessing the interacting effects of varying the thermal mean, variance, and autocorrelation in tandem (3.2); and measuring the repercussions of varying the autocorrelation of the observed thermal regimes experienced by 38 invertebrate species to compare how autocorrelation has modulated risk over time (3.3).

## 2 Methods

### 2.1 Model Description

To explore the relationship between temporal autocorrelation and thermal risk, we used logistic growth models to track population abundance through carefully varied thermal regimes by calculating population fitness (and corresponding changes in population size) at each time step according to that species’ TPC. We compared the effects of changing the mean, standard deviation, and autocorrelation of temperature on the thermal regime itself (Fig. 1, top left subfigures) and on the corresponding population dynamics (Fig. 1, bottom left subfigures) to establish the relationship between the ordering of temperatures and the risk of extinction, using spectral synthesis and spectral mimicry to precisely control the thermal distributions (Fig. 1, bottom right subfigures) and their temporal structure. We then compared how manipulating the autocorrelation of normally-distributed thermal regimes affected various extinction risk metrics for thermal generalist versus specialist populations, before testing the model on empirical TPCs by varying the autocorrelation of observed temperature time series from each organisms’ collection location.

**Figure 1:**
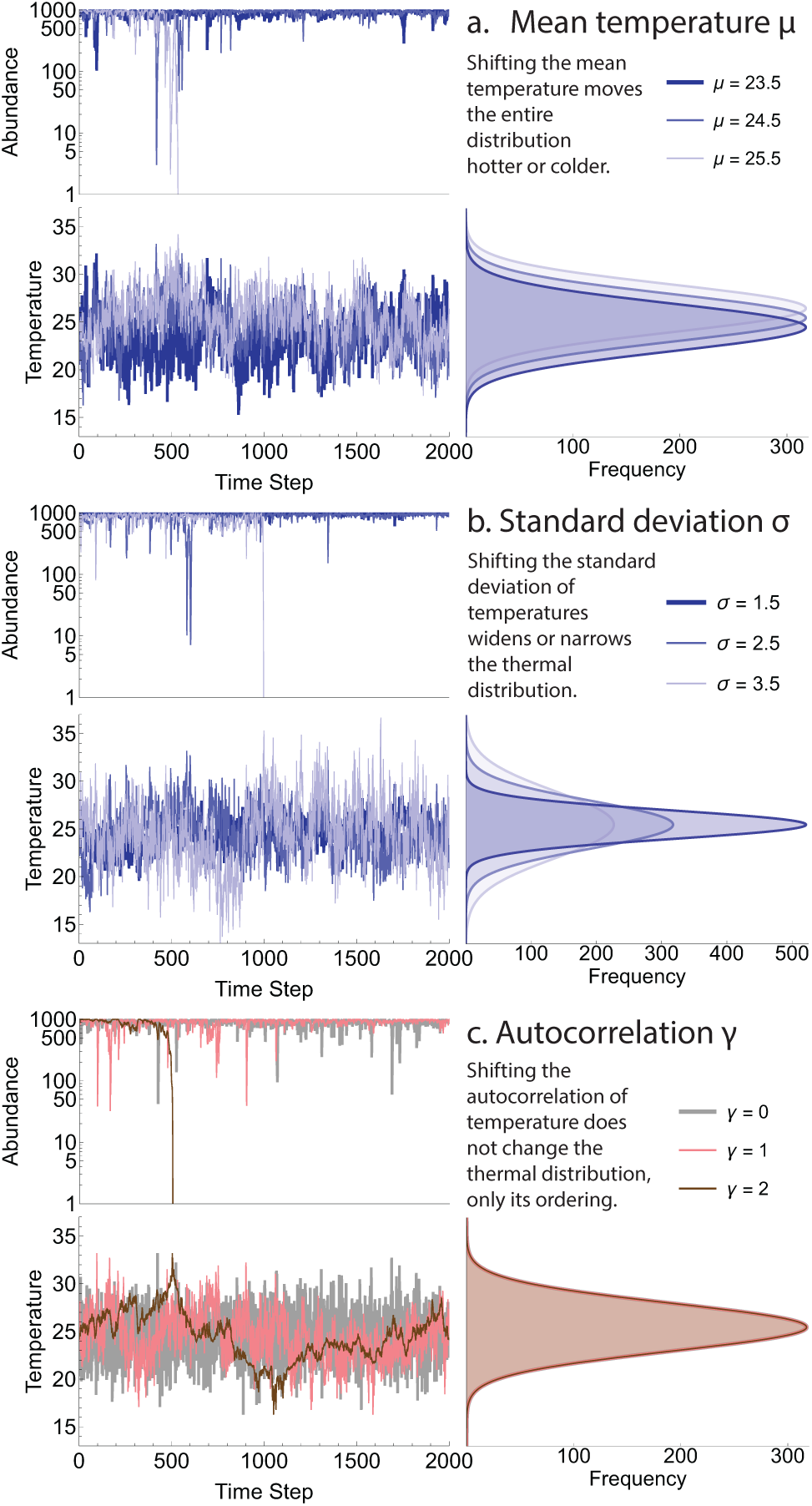
Example simulation runs with different transformations applied to each thermal regime. Sample population abundances and their corresponding thermal regimes are shown on the left, with overall thermal distributions for any simulation run on the right. Parameters not being varied are set to their mean values (*µ* = 24.5, *σ* = 2.5, *γ* = 1). Sample heat-driven extinctions occur around (a) *t* = 520 for *µ* = 25.5, (b) *t* = 1000 at *σ* = 25.5, and (c) *t* = 500 at *γ* = 2.

### 2.2 Population Dynamics

We incorporated thermal sensitivity into population dynamical models using Malthusian fitness, or the intrinsic population growth rate *r*. Defined as the difference between per-capita birth and death rates, this parameter has a well-established dependence on temperature (Amarasekare and Savage, 2012; Vasseur, 2020; Fey et al., 2021). Many studies use *r* as a proxy to describe the relationship between individual performance or population fitness and temperature (e.g., Deutsch et al., 2008; Kingsolver, 2009; Thomas et al., 2012); we took this one step further by incorporating the temperature-dependent *r* into the logistic growth model.

We calculated population sizes using the closed-form solution of the *r*-*α* logistic growth equation, assuming fitness *r* has temperature-dependence given by a TPC. We used the *r*-*α* formulation (Equation 1) of the logistic growth model (where *α* indicates the density-dependence) instead of the typical *r*-*K* formulation (Equation 2) (where *K* indicates the carrying capacity) due to the relative ease with which it handles negative fitness values (which importantly indicate population decline) and because it assumes a constant strength of density dependence *α* rather than a constant carrying capacity *K* (Long et al., 2007; Mallet, 2012; Vasseur, 2020); one can easily convert between the two by substituting *K* = *r/α*.

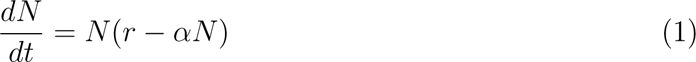

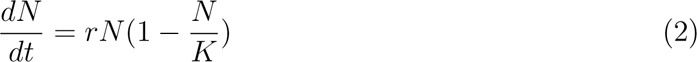

Because the logistic model declines asymptotically to 0 when *r <* 0, we defined an extinction threshold for our simulations as *N < K ×* 10*^−^*^6^, where *K* = 1*/α*. Shown results use *α* = 0.001, but changing the value of *α* impacts only the magnitude, not the behavior, of model dynamics (see Appendix S1).

### 2.3 Autocorrelated temperature variation

To determine the importance of incorporating temporally-explicit modeling into risk assessments, we compared populations experiencing the same thermal regimes but different amounts of autocorrelation. Temporally-blind metrics, or ones that do not take the ordering of temperatures into account, always predict the same level of risk for thermal regimes with the same statistical moments. Autocorrelation, however, may meaningfully impact the realized risk because increasing autocorrelation ‘clumps’ similar temperatures together within a series (Lawton, 1988; Vasseur and Yodzis, 2004; Schwager et al., 2006), leading to heatwaves or cold spells that could cause large differences between temporally-blind and temporally-explicit risk assessments.

To generate autocorrelated time series of temperature variation, we used a technique known as **spectral synthesis** (Cuddington and Yodzis, 1999; Halley et al., 2004; Fontaine and Gonzalez, 2005). This generates time series whose power spectra are characterized by a 1*/f^γ^* scaling property, where *f* is the frequency of the noise and 0 ≤ *γ* ≤ 2 is the spectral exponent (Halley, 1996; Vasseur and Yodzis, 2004). The spectral exponent characterizes the color of noise, where white noise (*γ* = 0) represents no autocorrelation and includes an equal mixture of all frequencies (such that fluctuations lack frequency dependence altogether), brown noise (*γ* = 2) represents high autocorrelation and has variance driven mostly by low frequencies (also known as a random walk or Brownian motion), and pink noise (*γ* = 1) lies in between (Halley, 1996; Vasseur, 2007). For a given spectral exponent *γ*, we generated a zero-mean normally-distributed time series *Y_γ_* from *t* = 1*, …, n* by summing *f* = 1*, …, n/*2 sine waves:

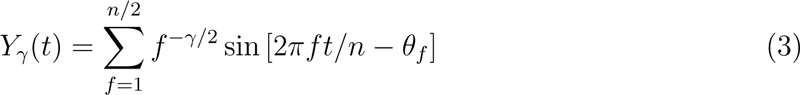

where *θ_f_* is a random vector of phase operators drawn from the uniform distribution [0, 2*π*) and the amplitude *f ^−γ/^*^2^ generates the power-law scaling (Vasseur, 2007). In practice, computing the sums in Equation 3 is inefficient and the same result can be obtained using an inverse fast Fourier-Transform (Cochran et al., 1967), which we implemented in our simulations using the function ‘InverseFourier’ in *Mathematica*.

To provide greater insights into the role of temporal autocorrelation on model dynamics, we used an algorithm known as **spectral mimicry** (Cohen et al., 1999) to apply the synthesized random temporally-autocorrelated sequence to a conserved set of temperature values. For each combination of temperatures with mean *µ* and standard deviation *σ*, we generated a normally distributed reference series *X_µ,σ_*of *n* temperatures by back-sampling values from the cumulative distribution function at *n* + 1 regularly-spaced intervals on (0, 1). This creates a ‘perfect’ distribution of temperature values that we permuted according to the ordinal ranking of values in *Y_γ_*. This permutation method imparts the desired autocorrelation structure in *Y_γ_* onto *X_µ,σ_* while preserving the same mean and variance (and, indeed, values) across different levels of autocorrelation. While the spectral exponent of *X_µ,σ_* can differ slightly from that of *Y_γ_*, the difference is generally very small and offset by the benefit of using the same values at each level of autocorrelation. Spectral mimicry thus ensures that comparisons between sets with the same temperature distribution but different spectral exponents reflect only differences in the ordering of the temperatures, not differences in the temperatures themselves.

### 2.4 Sample Thermal Performance Curves

Following Amarasekare and Savage (2012), we formulated TPCs as the difference between symmetric, unimodal birth/gain functions and exponential death/loss functions, such that *r* is defined by Equation 6. A Gaussian function, parameterized by the optimum growth temperature *TI* and a breadth parameter *β*, represents birth (Equation 4), while an exponential function, parameterized by a scaling factor *m_a_*, a rate factor *m_b_*, and an additive constant *m_c_*, represents death (Equation 5). This parameterization is used instead of the frequently employed piecewise Gaussian-quadratic (e.g., Deutsch et al., 2008; Vasseur et al., 2014; Duffy et al., 2022) because it is continuously differentiable, tied to the physiological reality that population fitness equals birth minus death rate, and allows populations to shrink at thermal extremes.

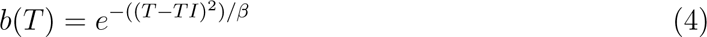

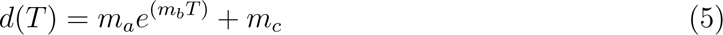

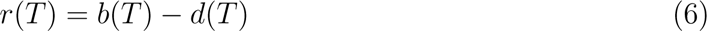

These equations produce the ubiquitous left-skewed TPC expected of most ectotherms (Angilletta, 2009; Dell et al., 2011; Rezende and Bozinovic, 2019). We characterized TPCs by four parameters: the thermal optimum *T*_opt_ (the temperature at which the the population exhibits peak growth), the thermal minimum *T*_min_ and maximum *T*_max_ (the *x*-intercepts beyond which death outpaces birth and the population shrinks), and the thermal breadth *T_b_* (equivalent to *T*_max_ − *T*_min_, the range of temperatures where fitness is positive and populations grow).

In initial models, we test two normalized TPCs: a temperate or generalist TPC, with a cooler thermal tolerance and wider thermal breadth, and a tropical or specialist TPC, with a warmer thermal tolerance but a narrower thermal breadth (Fig. 2). To emphasize the generality of our approach, neither curves’ parameters are linked to a specific study organism; however, both are similar to empirical parameters from the oft-used Frazier et al. (2006) dataset (see Appendix S2).

**Figure 2:**
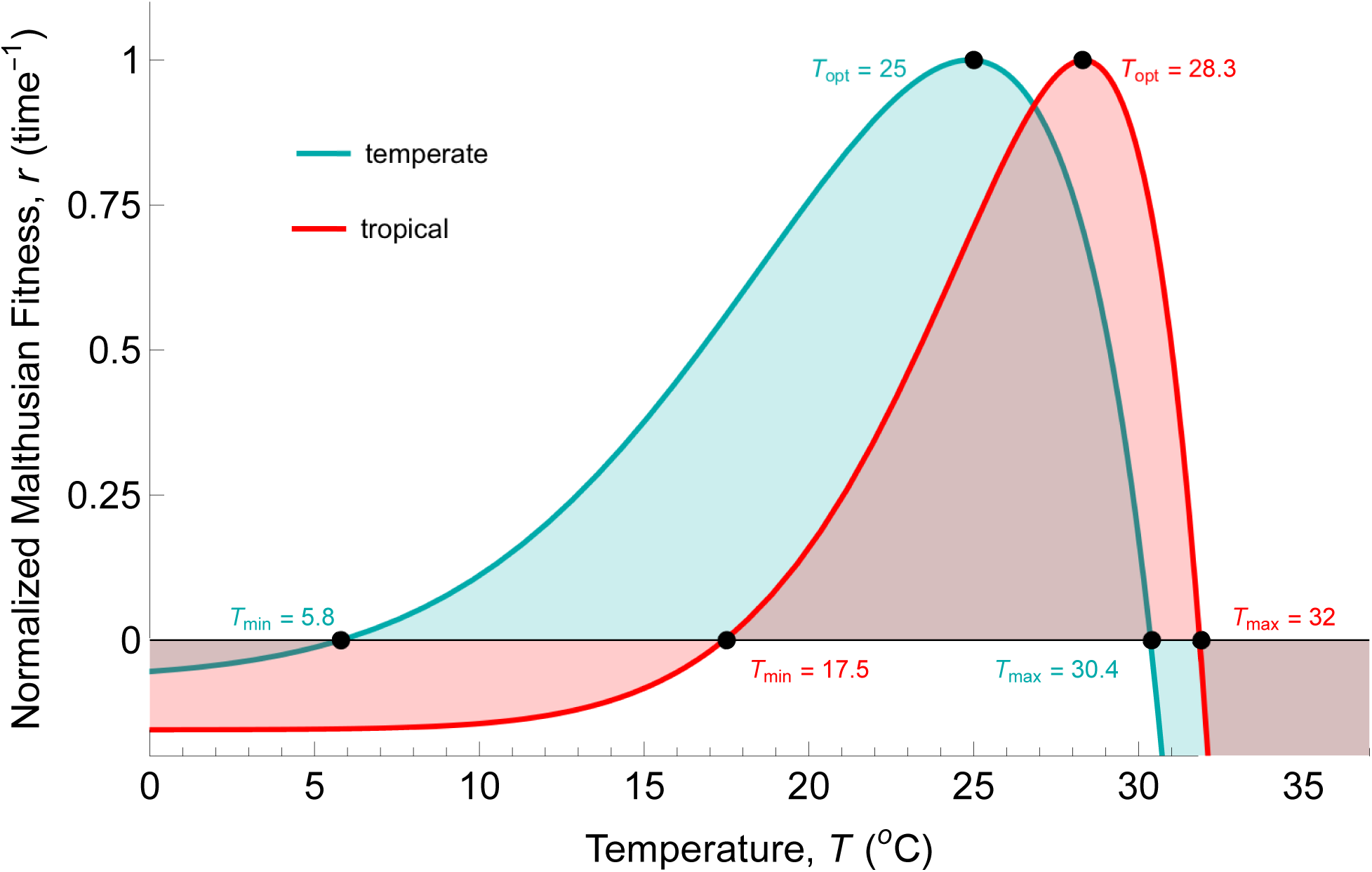
Temperate/generalist (1, teal) and tropical/specialist (2, red) TPCs. Model parameters for TPC 1: *TI* = 29, *β* = 180, *m_a_*= 5 × 10^−7^, *m_b_* = 0.4, and *m_c_* = 0.05; normalized by a factor of 0.755. Model parameters for TPC 2: *TI* = 35, *β* = 170, *m_a_* = 5 × 10^−7^, *m_b_* = 0.4, and *m_c_* = 0.05, shifted 2 degrees to the right relative to TPC 1; normalized by a factor of 0.4067.

### 2.5 Temporally-Blind Metrics

Before running simulations, we calculated several temporally-blind metrics, including the warming tolerance (WT), the difference between *T*_max_ and the mean environmental temperature, and the thermal safety margin (TSM), the difference between *T*_opt_ and the mean environmental temperature (Deutsch et al., 2008). Many other risk metrics rely instead on projected population size; for example, the IUCN Red List categorizes species as Vulnerable, Endangered, or Critically Endangered if they meet the risk criteria for any one of five categories (A. Population size reduction, B. Geographic range reduction, C. Small population size and decline, D. Very small or restricted population, or E. Quantitative Analysis) – four of which explicitly base categorizations on population size and one of which (B) is often a risk factor *because* of its impact on population sizes (IUCN, 2012). To calculate the projected population size in a temporally-blind fashion, we took the average fitness expected for a thermal regime *X_µ,σ_* divided by the density dependence parameter *α*.

### 2.6 Empirical TPCs and Climate Data

For our final model, we used the 38 species Frazier et al. (2006) global dataset of insect TPCs (gathered from 31 studies and 35 locations between 1974 and 2003) previously analyzed in Deutsch et al. (2008), Vasseur et al. (2014), and Duffy et al. (2022). For comparison, we fit this empirical data with the same Gaussian-quadratic TPC fit analyzed in earlier papers (see Appendix S2), allowing negative growth rates at high temperatures following Vasseur et al. (2014) and Duffy et al. (2022). To test the impact of changing autocorrelation levels on realistic thermal regimes – which are rarely perfectly normally distributed – we gathered a decade of observed historical (1994-01-01 to 2003-12-13) and recent (2014-01-01 to 2023-12-31) daily minimum and maximum temperatures from each location using Visual Crossing’s Weather Data Services (see Appendix S3).

We retained seasonal and diurnal fluctuations in permuted observational datasets to ensure that our spectral methods did not disrupt realistic and potentially relevant elements of the climate data. To maintain the relevance of seasonality but alter auto-correlation, we seasonally detrended the climate data before using spectral mimicry to manipulate its spectral exponent, then added the seasonal trend back in. This procedure allowed us to maintain the ‘power’ ratio of seasonal to non-seasonal components, the former of which was preserved and the latter of which had its autocorrelation altered. To maintain diurnal variation, we applied spectral mimicry to the mean of each day’s maximum and minimum temperatures, such that the days’ average temperatures are sorted with a specific autocorrelation level but the population experiences the corresponding extremes. We also estimated the spectral exponent for the unaltered climate data to use as a reference point for comparison. We then simulated the population dynamics of each species at each autocorrelation level for their respective thermal regimes and compared predicted extinction risks at the observed spectral exponents.

### 2.7 Simulations

All models were run in *Mathematica* V.13.0.1 for 1000 simulations of 2000 time steps for every autocorrelation level, thermal regime, and TPC. Because we incorporate a finite number of temperatures into each run, an appropriate time step size must be chosen to avoid obscuring results; relative to the generation time, overly long time steps cause population dynamics to average across all variation, while overly short ones result in population dynamics closely tracking all variation, regardless of the actual dynamics. As our TPCs were normalized with *r* ≤ 1, we used a time step size of 1 where neither averaging nor tracking dominates (see Appendix S4); we halved the time step size for the final model to accommodate the addition of 2 data points per day in the observational dataset. Each simulation began with an initial population size equal to the average population size expected for that thermal regime, such that *N*_0_ = *r̄*(*X_µ,σ_*)*/α*.

## 3 Results

### 3.1 Autocorrelation’s Effect on Abundance and Risk Metrics

For the first model, we recorded the population size at each time step for each normally-distributed thermal distribution and calculated the temporally-explicit risk metrics (mean population abundance, population stability, and proportion of extinct simulations) for two sample TPCs using four mean temperatures (*µ* = 23-26, intervals of 1), one standard deviation of temperature (*σ* = 3), and 21 autocorrelation levels (*γ* = 0-2, intervals of 0.1), then compared the calculated extinction risks with those found using temporally-blind metrics.

#### 3.1.1 Temporally-Blind Metrics

Given the mean temperatures, the temperate TPC (*T*_opt_ = 25, *T*_max_ = 30.4) had WTs of 7.4, 6.4, 5.4, and 4.4°C and TSMs of 2, 1, 0, and -1°C, respectively, at every level of variance and autocorrelation. The tropical TPC (*T*_opt_ = 28.3, *T*_max_ = 32) had WTs of 9, 8, 7, and 6°C and TSMs of 5.3, 4.3, 3.3, and 2.3°C, respectively, at every level of variance and autocorrelation. These WTs and TSMs indicate a trend of declining resilience with increasing temperatures, but suggest potential risk only for the temperate TPC and only at the hottest two temperatures. Projected population sizes (dotted lines, Fig. 3 (a), (d)) were all positive, with the highest population sizes projected successively from coolest to warmest means for the temperate TPC and from warmest to coolest means for the tropical TPC.

**Figure 3:**
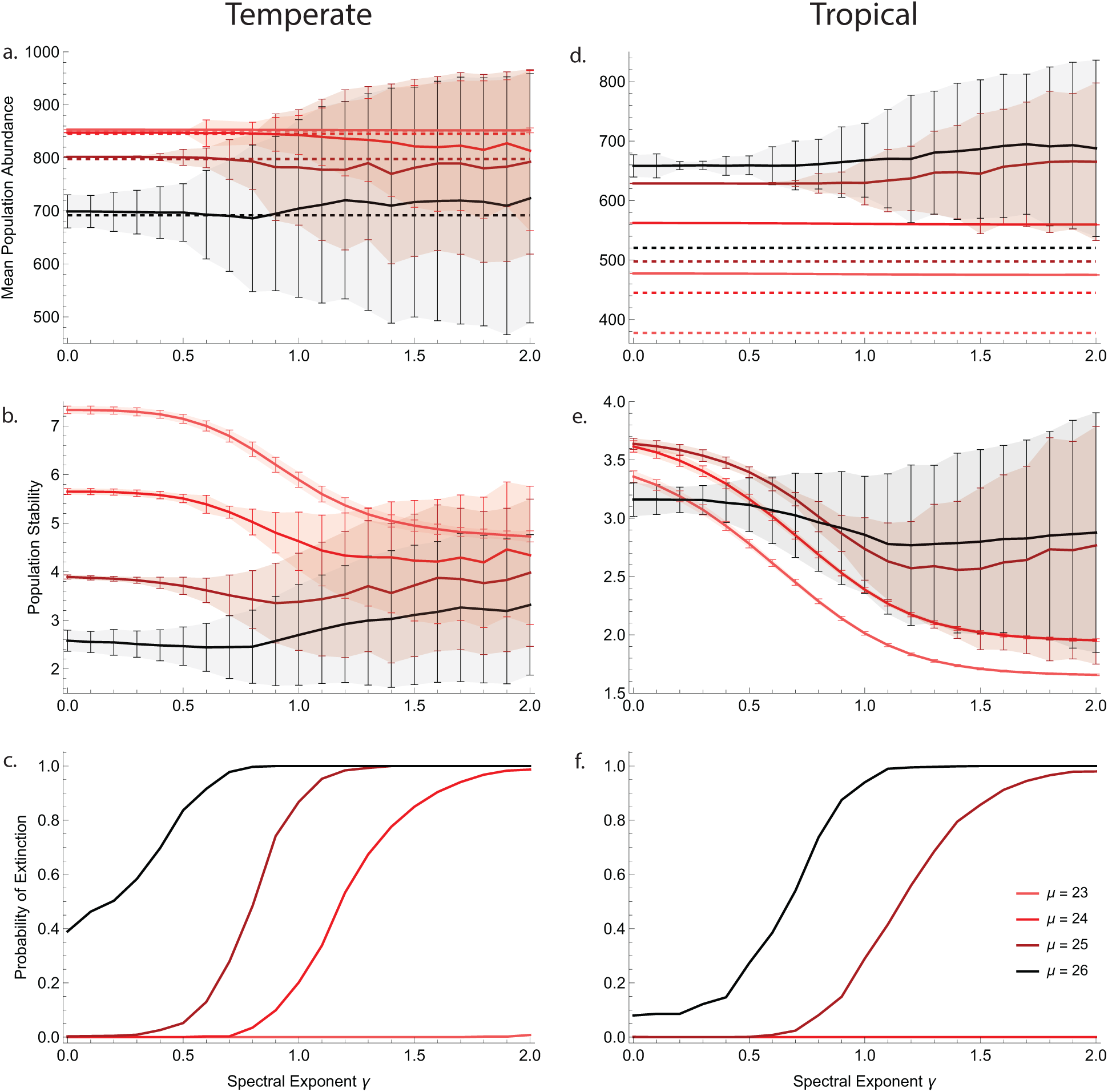
First model outputs, with the temperate/generalist TPC on the left and the tropical/specialist TPC on the right. Darker colors indicate higher mean temperatures. Error bars/shaded regions indicate the standard deviation of the given metric. Mean abundance plots (a, d) show the simulated mean abundance up to the time of extinction (solid lines) and the predicted mean abundance for the thermal regime *X_µ,σ_* (dotted lines). Stability plots (b, e) show the mean abundance divided by the standard deviation of abundance (inverse of the coefficient of variation; Duffy et al. (2022)) up to the time of extinction, with lower values indicating lower stability. Extinction plots (c, f) show the number of simulations out of the total where the population abundance dropped below the extinction threshold.

#### 3.1.2 Mean Population Abundance

The temporally-explicit model predicted the mean abundances shown in Fig. 3 (a) and (d) (solid lines). For the temperate TPC, population abundance declined as mean temperature increased. Each successive degree of warming resulted in comparatively greater declines and simulations closely matched the temporally-blind predictions. Autocorrelation affected mean abundance very little, although variability greatly increased at higher levels at hotter mean temperatures (growing error bars at higher *γ*s in Fig. 3 (a)). For the tropical TPC, population abundance increased with mean temperature. Each successive degrees of warming resulted in comparatively smaller increases and simulations greatly overshot the temporally-blind predictions. Again, autocorrelation had very little effect beyond increasing the variability of mean abundance for the hottest mean temperatures at *γ >* 0.5.

#### 3.1.3 Population Stability

For the temperate population, stability (defined as the mean population size divided by its standard deviation) had the expected negative correlation with increasing mean temperature (Fig. 3 (b)). The coolest mean temperatures (*µ* = 23, 24) allowed the most stable populations. Stability declined with increasing autocorrelation. At hotter mean temperatures (*µ* = 25, 26), the same trend occurred moving from white into pink noise (*γ <* 0.8), but stability then began increasing with autocorrelation level. This trend is primarily an artifact of the statistic itself: because the mean and standard deviation of abundance were calculated only *up until extinction*, the stability calculations included more runs at low autocorrelation levels (where very few runs went extinct) than at high autocorrelation levels (where many or most runs went extinct). The runs that persisted at high autocorrelation levels are, by nature, those with the highest, most ‘stable’ abundances, because the high levels of autocorrelation clumped the worst conditions of the thermal regime at the end of the run.

Overall, the tropical population was much less stable than the temperate one, but trends across mean temperatures differed (Fig. 3 (b)). Excluding the hottest thermal regime, stability increased with mean temperature at all autocorrelation levels; at the hottest mean temperature (*µ* = 26), stability was lower than every other regime under white noise conditions (*γ <* 0.3) but higher than every other under pink to brown noise conditions (0.9 < γ ≤ 2). Stability declined predictably with increasing autocorrelation across all mean temperatures, but the comparatively limited decline and slight uptick in stability under brown noise for the hottest two scenarios (*µ* = 25, 26) again demonstrated the ‘benefit’ of clumping stressful conditions late in the run in hotter regimes.

#### 3.1.4 Probability of Extinction

For both TPCs, the actual probability of extinction increased with both the mean temperature and the autocorrelation level (Fig. 3 (c) and (f)). The temperate population faced higher extinction risk than the tropical population across the board, though at the hottest two mean temperatures, both faced nearly a 100% chance of extinction at high levels of autocorrelation. While autocorrelation had no impact on extinction risk under completely permissive thermal regimes (*µ* = 23 for temperate, *µ* = 23, 24 for tropical), it had a pronounced impact on extinction risk under more stressful ones; for example, at *µ* = 25, the temperate species shifts from an extinction risk near 0% to an extinction risk near 100% in the range of 0.5 *< γ <* 1.

#### 3.1.5 Comparing Metrics

Overall, population abundance was a poor predictor of extinction risk: all mean temperatures resulted in simulated mean abundances far above the extinction threshold, and, indeed, for the tropical population, the lowest mean abundances (*µ* = 23, 24) corresponded with the lowest extinction risk. The predicted temporally-blind abundance indicated even higher population sizes and no extinction risks at all. Population stability was a fair predictor of *relative* extinction risk, particularly for the temperate TPC and at lower levels of autocorrelation (e.g., declining stability from 0 *< γ <* 1 corresponding with increasing temperate extinction risk); however, it was an unreliable predictor of risk for the tropical TPC (e.g., higher stability at *µ* = 26 than *µ* = 23, 24, 25 for *γ >* 0.9, despite far higher extinction risk for the former) and sometimes at higher levels of autocorrelation (e.g., increasing temperate stability for *γ >* 1, despite high extinction risk).

### 3.2 Extinction Risk Across the Thermal Landscape

For the second model, we studied the probability of extinction across a broader parameter space by calculating the proportion of extinct simulations at each of 7 mean temperatures (*µ* = 23-26, intervals of 0.5), 7 standard deviations (*σ* = 1-4, intervals of 0.5), and 21 autocorrelation levels (*γ* = 0-2, intervals of 0.1). Collecting only extinction time (not sequential population sizes) allowed us to test a broader parameter set, including the impacts of widening or narrowing the range of the thermal regime by changing the standard deviation (see Fig. 1 (b)). Due to their similarity, only results for the temperate TPC are shown in the main text; tropical results are shown in Appendix S5.

#### 3.2.1 Proportion Extinct

Fig. 4 shows the proportion of extinct runs for every combination of parameters; blue indicates no extinction events, while red indicates extinction in every run. The initial model’s results occur for the parameter set shown in Fig. 4 (a), subplot 5, with *σ* = 3. As expected, low parameter values – permissive thermal regimes with low variability and white noise coloration – resulted in no extinction risk, and increasing any individual parameter increased risk (trends from blue to red moving down and right across each sub-plot). However, the relevant magnitude of change varied between parameters. Changing the level of autocorrelation, for example, had no effect when the standard deviation is too low (*σ <* 2) or too high (*σ >* 3.5); in those cases, the overall variation is either so low that extinctions cannot be generated regardless of the temperatures’ ordering or so high that even randomly scattering the extremes throughout the simulation cannot save the population. Increasing autocorrelation had the most pronounced impacts on extinction risk in middling thermal regimes (e.g., Fig. 4 (a), 2 ≤ σ ≤ 3.5).

**Figure 4:**
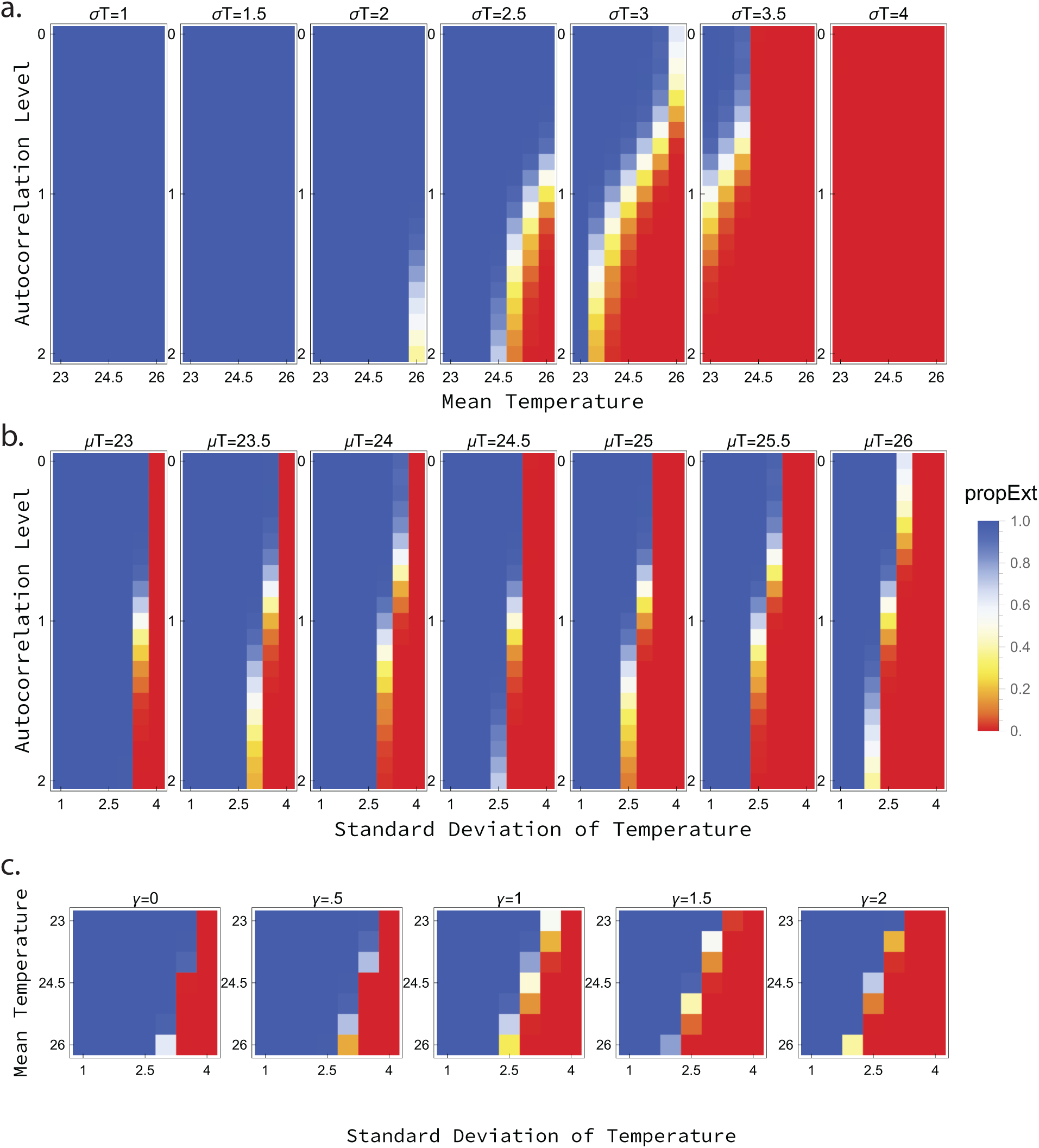
The proportion of extinct runs out of 2000 at each autocorrelation level and thermal regime for the temperate TPC. (a) Mean temperature against autocorrelation level at each standard deviation. (b) Standard deviation against autocorrelation level at each mean temperature. (c) Standard deviation against mean temperature at each autocorrelation level.

#### 3.2.2 Time to Extinction

Fig. 5 shows the mean time to extinction across all runs; these outputs are the same model results as shown in Fig. 4, but now shaded by how early the populations went extinct. Blue again indicates no extinctions (equivalent on the color bar to “extTime = 2000”). All other colors indicate that some proportion of the runs went extinct, with yellow representing later average extinction times and red representing earlier average extinction times.

**Figure 5:**
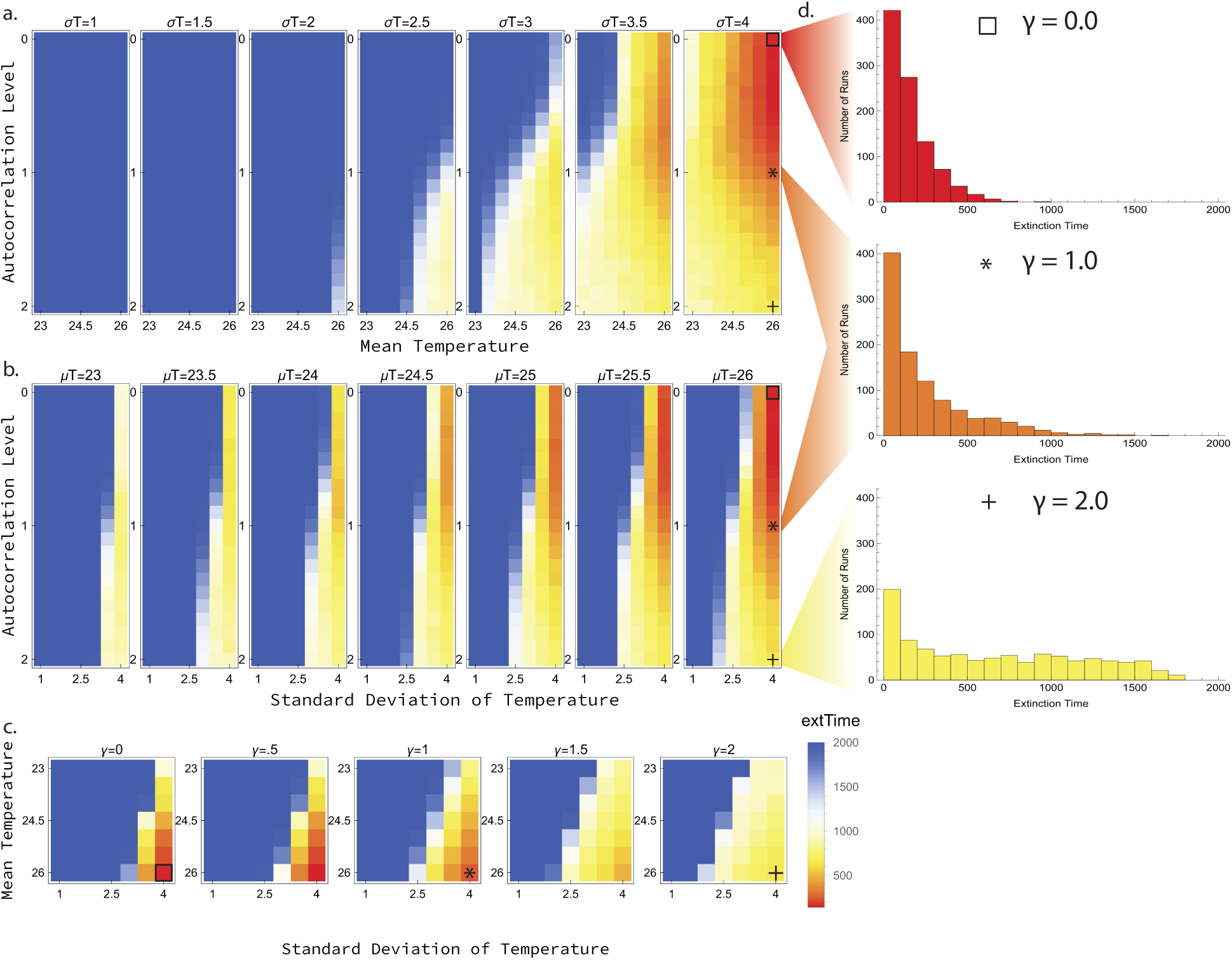
The average extinction time across runs for the temperate TPC at each autocorrelation level and thermal regime. (a) Mean temperature against autocorrelation level at each standard deviation. (b) Standard deviation against autocorrelation level at each mean temperature. (c) Standard deviation against mean temperature at each autocorrelation level. (d) Histograms of the extinction times at the highest mean and standard deviations of temperatures at different autocorrelation levels. Squares (*γ* = 0), stars (*γ* = 1), and plus signs (*γ* = 2) indicate the location of the histograms in each set of heat plots.

While increasing autocorrelation resulted in a higher *proportion* of extinctions (less blue moving down subplots in Fig. 5 (a), (b) and across subplots in (c)), it did not uniformly result in *earlier* average extinction times. Instead, low levels of autocorrelation corresponded with early extinctions under stressful thermal regimes; for example, when *µ* = 26 and *σ* = 4, the extinction times are distributed as shown in the histograms of Fig. 5 (d). For the white noise scenario (*γ* = 0, top red histogram), most populations went extinct before the 500^th^ time step. In contrast, for the brown noise scenario (*γ* = 2, bottom yellow histogram), many populations persisted past the 1500^th^ time step – even though just as many total extinctions occurred. This paradox is due to the extremity of the thermal regimes; when the combination of hotter means and larger standard deviations pushes the extreme temperatures hot enough, the corresponding population decline is so precipitous that scattering each individual extreme temperature randomly throughout the simulation (white noise) is riskier than clumping them all together (brown noise), because any extreme temperature on its own could cause an extinction. In other words, individual heat events are so deadly that high autocorrelation lengthens persistence times simply by clumping the extremes together later in the simulation run.

### 3.3 Impacts of Autocorrelation on Global Risk Assessments

For the final model, we compared a decade of historical and recent observed temperature time series (e.g., Fig. 6 (a)) across 21 autocorrelation levels (*γ* = 0-2, intervals of 0.1) using empirical TPCs from 38 insects. In contrast to initial models, the thermal regime’s mean and standard deviation are not artificially varied; we checked only how autocorrelation mediated risk in historical thermal regimes compare to recent ones, benchmarking simulations with the estimated spectral exponent observed in the temperature time series.

**Figure 6:**
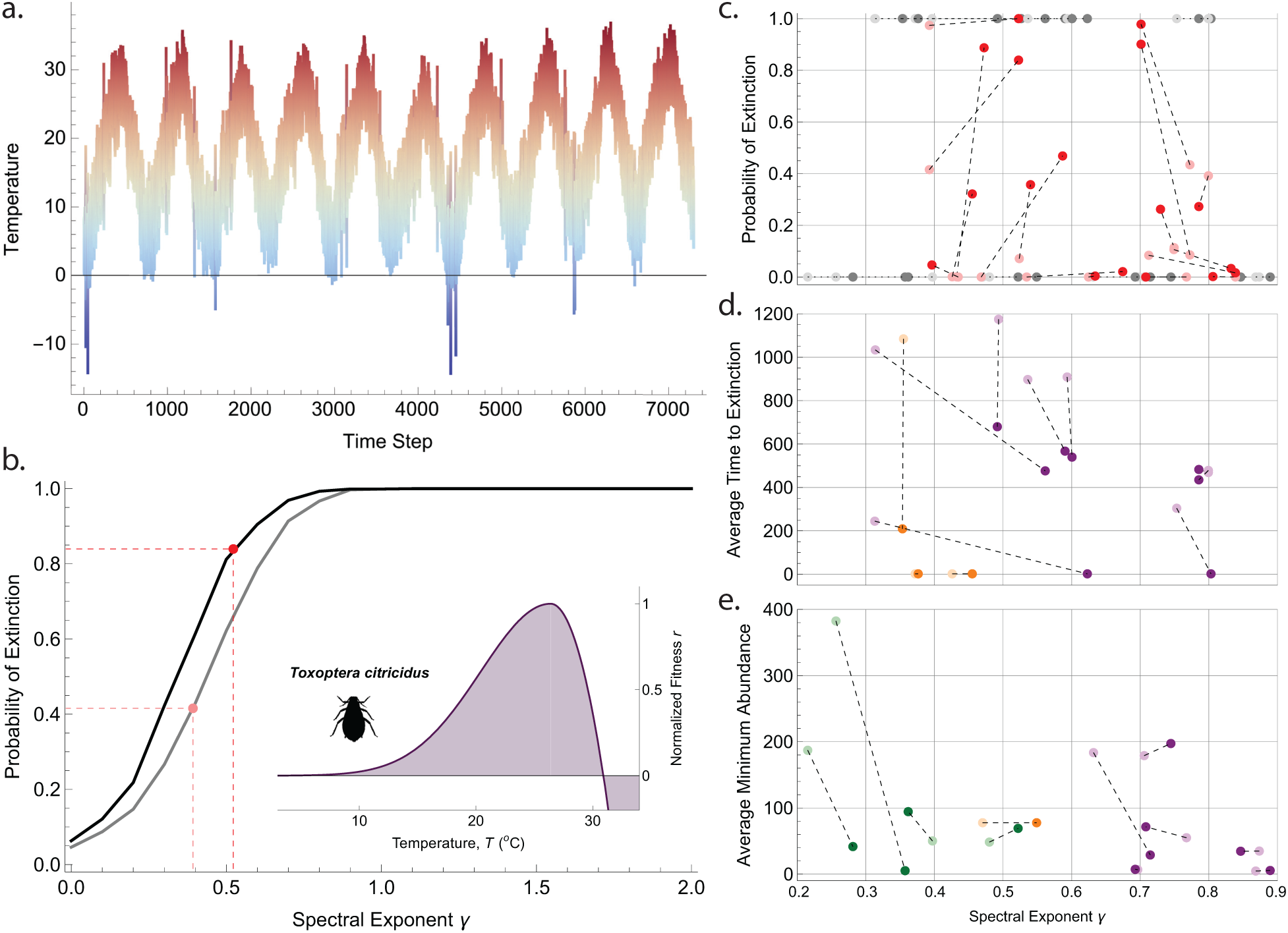
(a) Example temperature time series of the observed historical (1994-2003) temperatures at 34.98*^o^*N, 138.4*^o^*W, the collection location of *Toxoptera citricidus*, re-sorted with an autocorrelation level of *γ* = 1 with seasonal and diurnal fluctuations included. (b) Proportion of runs where *T. citricidus* went extinct at each autocorrelation level given observed historical (gray line, 1994-2003) and recent (black line, 2014-2023) temperatures and its empirical TPC (purple subplot). Actual autocorrelation levels of the observed temperature time series, and their corresponding extinction risks, are marked in pink (historical) and red (recent). (c) Expected probability of extinction given observed spectral exponents for each of the 38 species; historical (pink) and recent (red) points for any given species are connected with a line. Gray points indicate species that always (top) or never (bottom) went extinct. (d) Average time to extinction for the 11 species that always went extinct. (e) Average minimum abundance for the 11 species that never went extinct. For the final plots, species are colored by geographic region (purple for Northern Hemisphere, green for tropics, orange for Southern Hemisphere), with the recent time period always indicated by darker shading.

#### 3.3.1 Risk responds to autocorrelation under realistic conditions

Using real thermal distributions and including the observed diurnal and seasonal variation, we still found the same S-shaped relationship between extinction risk and spectral exponent observed with the sample thermal performance curves and normally distributed temperatures (Fig. 6 (b)). The fluctuations buffered risk somewhat by forcing nightly or seasonal relief from stressful temperatures, but we could recover the same S-shaped curve for every species by varying the extinction thresholds (for example, given a TPC, increasing the extinction threshold from *K* × 10^−6^ individuals to *K* × 10^−3^ for more permissive thermal regimes or decreasing it to *K* ×10^−9^ for more stressful ones). The remainder of the analysis was performed with an extinction threshold of 1 individual.

#### 3.3.2 Projected species-level impacts

The output for each species incorporates three results: whether spectral exponents increased or decreased over time, whether background thermal regimes made extinction more or less likely across all autocorrelation levels, and how the probability of extinction changed based on observed spectral exponents. Full results for every species, including TPCs, histograms of historic and recent temperatures, and extinction curves, are found in Appendix S6: Fig. S1.

The spectral exponent increased at 23 of the 34 collection locations (67.6%). Out of the 38 species tested, the recent thermal regime increased extinction risk above historic levels for 16 (42.11%), decreased extinction risk for 4 (10.53%), and did not affect extinction risk for the remaining 20 (47.37%). At observed autocorrelation levels from the empirical datasets (Fig. 6 (c)), extinction risk increased for 13 species (34.21%), decreased for 3 species (7.89%), and remained unchanged for 22 species (57.89%); of those 22, 11 went extinct in every simulation while 11 never went extinct at all (see below).

Regardless of whether the spectral exponent itself increased or decreased between time periods, extinction risk increased exclusively in scenarios that became more sensitive to autocorrelation (extinction curves shifting left; e.g., Fig. 6 (b)).

To assess whether risk increased for those species whose probability of extinction was 100% in both historic and recent simulations, we looked at the mean time to extinction across simulations at each autocorrelation level (Fig. 6 (d); full results in Appendix S7: Fig. S1). As expected from earlier modeling results, high levels of autocorrelation incrementally increased persistence time by clumping stressful conditions later in runs (see 3.2.2). However, at the measured spectral exponents, populations went extinct faster under recent conditions in 8 out of 11 scenarios (average decrease of 57.71%, or 201 days); one species went extinct slower (increase of 2.99%, or 7 days); and two species always went extinct immediately, indicating that the regimes were too stressful for persistence under any circumstance.

For species whose probability of extinction was 0% under both historic and recent temperatures, we looked instead at the minimum population size encountered across simulations at each autocorrelation level (Fig. 6 (e); full results in Appendix S7: Fig. S2). We saw a distinct trend of declining minimum abundance with increasing spectral exponent in all cases. At the measured spectral exponents, minimum population sizes were smaller under recent temperatures for 5 out of 11 species, but the difference between regimes where abundance increased were general very small (average increase of 16 individuals) compared to those were abundance decreased (average decrease of 136 individuals).

In summary, 26 species faced higher risk under recent thermal regimes (68.42%) compared to 10 species with lower risk (26.32%) and 2 species with no change (5.26%). 14 species faced projected extinction risk over 90% under recent temperature scenarios. Geographically, increased versus decreased risk was divided 60%/40% in the tropics (5 species), 68%/32% in the Northern Hemisphere (25 species), and 75%/0% in the Southern Hemisphere (8 species, with the remaining 25% facing 100% extinction risk across scenarios). The level of temporal autocorrelation affected extinction risk for 92% of tested species.

## 4 Discussion

Our models show that the impact of changing thermal regimes on population persistence depends on the combination of parameters: increasing means and extremes do accelerate extinction risks, but as conditions grow more stressful, that effect is increasingly mediated by temporal autocorrelation. Intensifying heatwave regimes vastly reduce predictability, stability, and persistence probability; those impacts grow more pronounced as the background thermal regime becomes more stressful. Indeed, while increasing a thermal regime’s variance causes the pronounced increase in extinction risk expected from earlier results (e.g., Vasseur et al., 2014; Slein et al., 2023), this effect does not drive extinction risk alone unless the scenario is extremely permissive or extremely stressful. We further found that when experiencing a thermal regime stressful enough to generate extinctions but not stressful enough to guarantee them, realistic autocorrelation values often fell along the steep slope between probable persistence and probable extinction. While increasing means and variances of temperatures shape the background thermal landscape, the extent of autocorrelation fundamentally determines whether realistic thermal regimes are survivable or deadly.

Autocorrelation has previously been shown to increase extinction risk of populations in models where the environment additively changes the population growth rate (Petchey et al., 1997) or where it alters the carrying capacity of a population (Cuddington and Yodzis, 1999). Schwager et al. (2006) clarified that whether red noise increases or decreases extinction risk over white noise depends upon the number of catastrophic events: redder noise decreases extinction risk when its ‘redness’ ensures high similarity between values and thus reduces the likelihood of catastrophes. This distinction, while true, conflates the impact of higher autocorrelation with the impact of lower variance; our use of spectral mimicry has allowed us to show that when variance is held constant, red noise unambiguously increases extinction risk over white noise.

Here, we have incorporated temperature variation through a well-established and mechanistic non-linear relationship using TPCs, the shape of which incorporates the stressful impacts of heat events via the accelerating decline of the TPC at high temperatures (Fig. 2). Autocorrelation is known to increase ‘tracking’ – the relationship between the actual population size and the equilibrium at any given temperature (Vinton and Vasseur, 2020) – because, for a given variance, changes at short timescales are smaller, allowing the population to more closely follow its equilibrium. When temperatures enter the stressful range in this model, *r*(*T* ) *<* 0 and the equilibrium population size is 0; the relatively smaller variations at short timescales act to increase the length of stressful periods, increasing the likelihood that a population will reach critically low sizes and cross the extinction threshold. The ability of a population to ‘track’ the equilibrium in our model depends on the magnitude of *r*(*T* ); as heat stress leads to more negative values of *r*(*T* ), the likelihood of extinction thus increases more quickly.

Standard risk metrics, like TSMs and population abundances, become significantly worse predictors of persistence under these higher autocorrelation scenarios. Comparing predicted outcomes for a generalist, temperate-adapted species and a specialist, tropical-adapted species showed that hotter thermal regimes can increase specialist abundance and extinction risk simultaneously. It will be critical for scientists and policymakers alike to account for this aspect of climate change under warming scenarios with increasing autocorrelation, and may be particularly important in systems already known to experience higher levels, like marine environments (e.g., Vasseur and Yodzis, 2004; Duffy et al., 2022).

While this model provides the most comprehensive analysis of the interactions between changing thermal regimes, population dynamics, and population persistence to date, it includes a considerable number of assumptions. By choosing logistic growth of a density-dependent population, we have incorporated the built-in assumptions of that model, including that the thermal response of carrying capacity *K* will be the same as that for fitness *r* (Bieg and Vasseur, 2024). We have ignored ecological interactions (Dell et al., 2014; Fey and Vasseur, 2016) and assumed that empirical TPCs are a good proxy for population growth rates (Sinclair et al., 2016). Perhaps most critically, we have assumed that those TPCs are static across space and time: each individual in these models has the same thermal tolerance regardless of life stage (Kingsolver et al., 2011), thermal history (Rebolledo et al., 2021), population structure (Angert et al., 2011), and body state (Brett et al., 1969; Huey and Kingsolver, 2019). We have not allowed organisms to evolve over time (Forester et al., 2022), experience acclimation (Phillips et al., 2016), adaptation (O’Donnell et al., 2018), and heat shock (Boopathy et al., 2022), or to physiologically and behaviorally thermoregulate to decouple their body temperature from the air temperature (Kearney et al., 2009; Sears et al., 2016). We have further – and once again – extrapolated conclusions from a dataset of 38 invertebrates that has been repeatedly analyzed over the years (Deutsch et al., 2008; Vasseur et al., 2014; Duffy et al., 2022), despite its biased focus on 50+ year old lab populations of economically important pest insects spread unevenly across the globe (Frazier et al., 2006). While these various assumptions may each worsen or improve the predictions we have made, the general patterns of how autocorrelation mediates extinction risk will hold. Exploring whether more realistic assumptions about how organisms interact with their thermal environments exacerbate or negate these risks should be an area of critical consideration moving forward.

We have shown that given a thermal regime, extinction risk can shift dramatically due solely to changes in the temporal autocorrelation – even when classic metrics like average population size and TSMs indicate no risk. While the statistical quantities describing a regime’s temperature distribution provide valuable information about the prevalence of stressful events when paired with an organism’s TPC, the picture is not complete without consideration of the order in which those temperatures occur. It is thus critical to consider the noise structure of stressful thermal regimes when assessing extinction risks, especially as temperatures grow more stressful and heatwave regimes continue to intensify.

## Acknowledgements

Funding for this study was provided by Yale University.

## Author Contributions

AR and DV conceptualized the study and model design, AR gathered data, performed analyses, and wrote the first draft, and both authors contributed to revisions.

