## Appendices for "Order matters: Autocorrelation of temperature dictates extinction risk in populations with nonlinear thermal performance"

**Appendix S1** *Sensitivity to density dependence*

We calculated population dynamics using the $r-\alpha$ formulation of the logistic growth model, which incorporates the density dependence parameter $\alpha$. Model results throughout were calculated with $\alpha = 0.001$ and, consequently, a carrying capacity of $K = 1000$. However, changing the value of $\alpha$ affects only the magnitude of the behavior observed, not the behavior itself, as shown in Fig. S1; the population stability and probability of extinction are nearly identical at the different $\alpha$ values, but the population abundance scales down according to the relationship between $\alpha$ and $K$. Notably, we calculate the extinction threshold $N_{ext}$ to scale proportionally with density dependence: $N_{ext} < K \times 10-6 = 1/\alpha \times 10-6$.

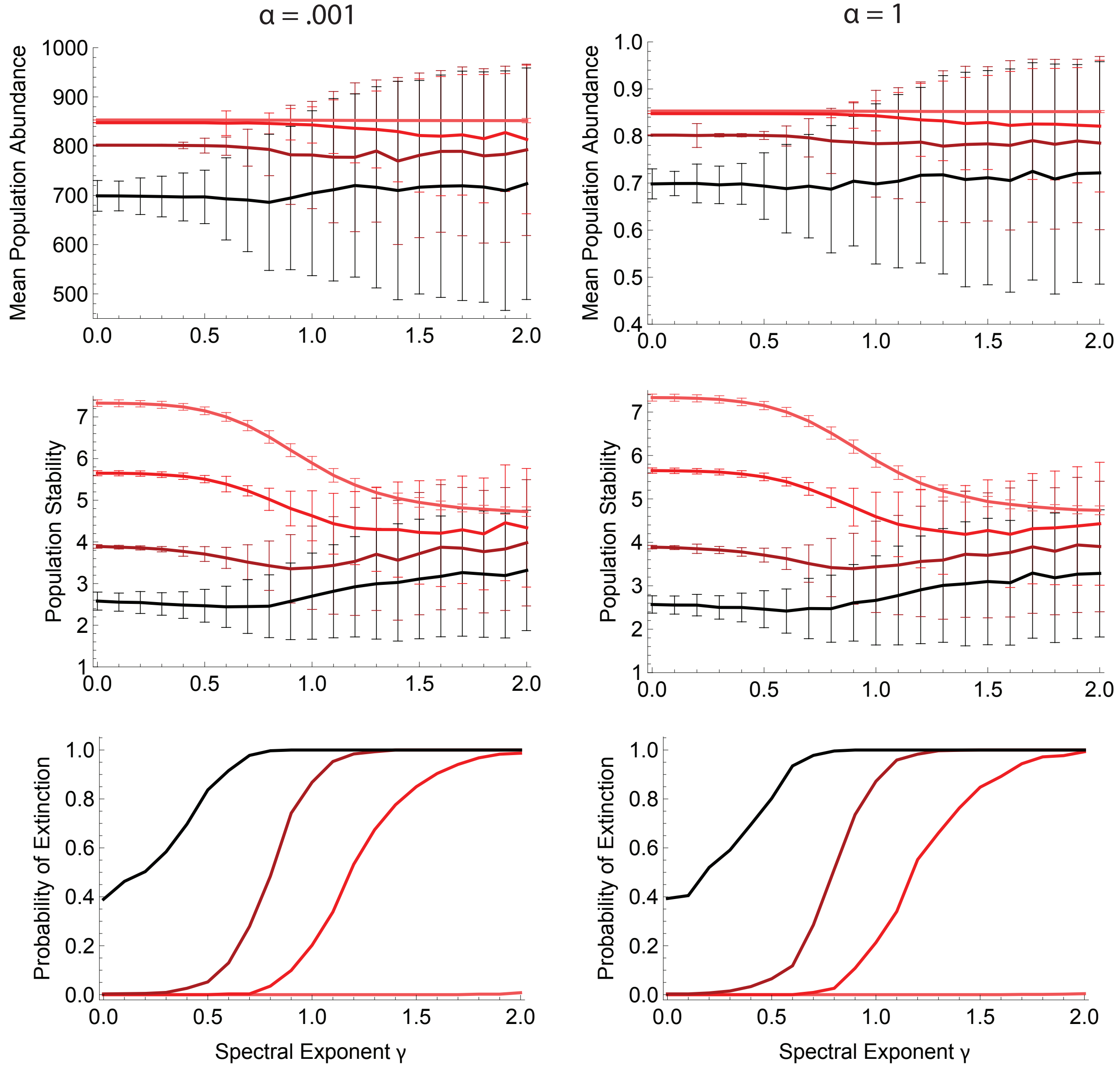

**Figure S1:** Example of the insensitivity of model results to a four-degree magnitude change of $\alpha$, utilizing the first model and the temperate TPC. On the left, the model results shown in Fig. 3 a, b, and c ($\alpha = 0.001$, $K = 1000$). On the right, the model results for the exact same model run with different density dependence ($\alpha = 1$, $K = 1$). The same trend holds for any value of $\alpha$.

**Appendix S2** *Basis of Phenomenological TPCs*

For initial models, we used example temperate and tropical TPCs similar to those in the dataset of 65 generalist crop-pest invertebrates originated in Frazier et al. (2006) and subsequently used – in an abbreviated form of 38 of the original species – in the analyses of Deutsch et al. (2008), Vasseur et al. (2014), and Duffy et al. (2022). Empirically determined parameters for the five species most similar to each phenomenological curve are shown in Table S1, with similarity determined by mean absolute error (MAE). In Fig. S1, we compare the TPCs with the phenomenological parameters (colored lines) to those with the empirical parameters (gray lines). For this comparison, we use the Gaussian-Quadratic TPC fit employed by Deutsch et al. (2008):

$$r(T) = \begin{cases} \exp\left(-\left(\dfrac{T - T_{\mathrm{opt}}}{2\sigma_p}\right)^2\right) & \text{for } T \leq T_{\mathrm{opt}} \\ 1 - \left(\dfrac{T - T_{\mathrm{opt}}}{T_{\mathrm{opt}} - T_{\mathrm{max}}}\right)^2 & \text{for } T > T_{\mathrm{opt}} \end{cases} \qquad \text{(S1)}$$

where $\sigma_p = (T_{\mathrm{opt}} - T_{\mathrm{min}})/4$. This is not the same set of equations used in the analysis of initial models (see 2.4), but is employed here simply to illustrate the similarities between parameters; the primary discernible difference between the Gaussian-Quadratic fit and our own is that their TPC is bounded above 0 in the original formulation, whereas we allow it to go negative below $T_{\mathrm{min}}$ and above $T_{\mathrm{max}}$. We do use the Gaussian-Quadratic in the final model to allow comparisons with previous studies; the full set of parameters for each species can be found in speciesparams.csv.

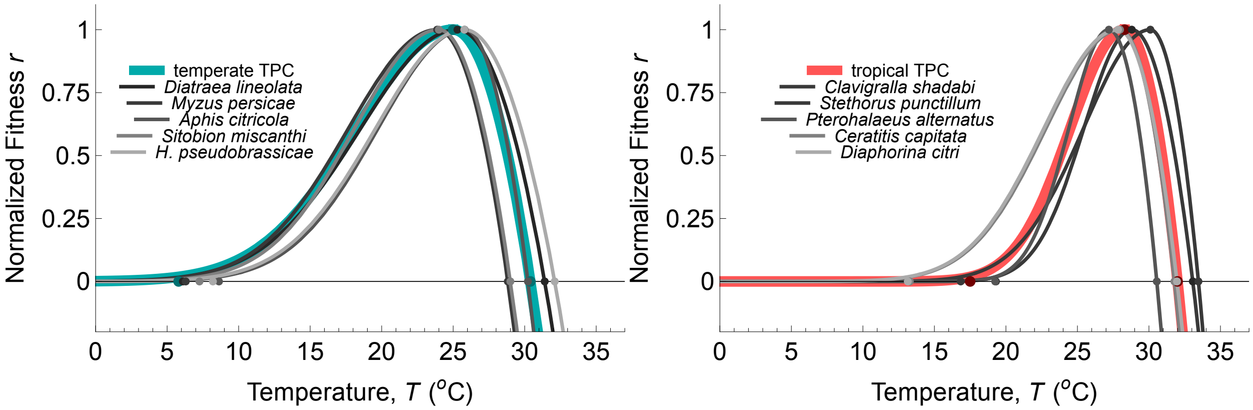

**Figure S1:** Comparison between phenomenological TPCs and empirical TPCs, with the used TPCs in color and empirical insect TPCs in gray. All displayed TPCs are drawn with the Gaussian-Quadratic fit, instead of the birth minus death framework (2.4), for the sake of comparison. Dots show location of $T_{\mathrm{opt}}$, $T_{\mathrm{max}}$, and $T_{\mathrm{min}}$ for each TPC.

**Table S1:** Parameters for the 5 species most similar to each phenomenological TPC.

| Species | Collection Location | $T_{opt}$ | $T_{max}$ | $T_{min}$ | MAE |
|---|---|---|---|---|---|
| *Temperate Comparisons* | | | | | |
| *Diatraea lineolata*<br>Neotropical Cornstalk Borer | Río Bravo, Mexico | 25.3 | 31.4 | 6.1 | 0.533 |
| *Myzus persicae*<br>Peach Aphid | Columbia, MO, USA | 23.88 | 28.82 | 6.3 | 1.067 |
| *Aphis citricola*<br>Citrus Aphid | Shizuoka, Japan | 25.8 | 30.23 | 8.6 | 1.267 |
| *Sitobion miscanthi*<br>Indian Grain Aphid | Sydney, NSW, Australia | 24 | 29 | 7.26 | 1.287 |
| *Hyadaphis pseudobrassicae*<br>Turnip Aphid | Columbia, MO, USA | 25.78 | 32.1 | 8.2 | 1.627 |
| *Tropical Comparisons* | | | | | |
| *Clavigralla shadabi*<br>African Pod Bug | Abomey Calavi, Benin | 28.8 | 33.1 | 19.3 | 1.123 |
| *Stehorus punctillum*<br>Spider Mite Destroyer | Quebec City, Canada | 30.1 | 33.46 | 16.84 | 1.307 |
| *Pterohalaeus alternatus*<br>Australian Wire Worm | Boonarga, QLD, Australia | 27.2 | 30.6 | 19.3 | 1.433 |
| *Ceratitis capitat*<br>Mediterranean Fruit Fly | Maui, HI, USA | 27.98 | 31.78 | 13.28 | 1.587 |
| *Diaphorina citri*<br>Asian Citris Psyllid | Pompano Beach, FL, USA | 27.84 | 31.96 | 13.14 | 1.620 |

**Appendix S3** *Climate Data*

We obtained climate data from Visual Crossing's data service, Visual Crossing Weather (https://www.visualcrossing.com/), using their Weather Query Builder interface. To obtain daily minimum and maximum temperatures, searches were run on the geographic collection coordinates of each species and the selected time ranges (historical: 01-01-1994 to 12-31-2003; present: 01-01-2014 to 12-31-2023) in metric data units with outputs 'datetime', 'tempmax', 'tempmin', and 'source'. The "Data Selection" option allowed us to choose the included data sources, indicated by the final output. We used "Historical station observations" (indicated by 'obs') or, in the case of missing observation data, "Statistics" (indicated by 'stat') run by the company to estimate likely values; queries occasionally resulted in blank rows of data attributed to a "Combined" source (indicated by 'comb'), in which case the query was run again on the missing days with Data Selection of only "Statistics" and manually filled in. Overall, there were 792 statistically generated points across the entire dataset; these points, spread across 14 of the 34 locations and all from the historical time range, represented 0.638% of the historical dataset and 0.319% of the total dataset. Once datasets were downloaded and cleaned, we removed February 29 of each included leap year (historical: 1996, 2000; present: 2016, 2020) because our function for varying autocorrelation (2.6) while maintaining seasonal and diurnal variation requires an equal number of points in every year; this accounted for 2 points per query, or 0.055% of the total dataset (Visual Crossing Corporation, 2024).

**Appendix S4** *Time Step Size*

Use of a discrete model necessitates a choice of time step size $\delta t$. If the choice of $\delta t$ is too short, the population dynamics will closely track the average, unchanging carrying capacity $K$ (e.g., yellow line in Fig. S1) dictated by the average fitness value for the temperature regime $Z_i$ (Fig. S1, dotted black line):

$$K = \frac{\left(\frac{\sum_{1}^{t_{\max}} r(Z_i)}{t_{\max}}\right)}{\alpha} \tag{S1}$$

Such short time steps might be analogous, for example, to the population experiencing a stressful temperature for only one second, which would not have any great impact on birth or death rates.

If the choice of $\delta t$ is too long, however, the population dynamics will be impacted too significantly by every temperature (e.g., red lines in Fig. S1) and, instead of converging to the average carrying capacity, will closely track the time-varying equilibrium $K_i$ calculated at every single temperature (Fig. S1, dashed black line):

$$K_i = \frac{r(Z_i)}{\alpha} \tag{S2}$$

This will result in extreme fluctuations of population sizes, including detrimental crashes every time $r(Zi) < 0$. Such long time steps might be analogous to the population experiencing every stressful temperature for an entire week, which would skyrocket death rates and decimate birth rates no matter how permissive adjacent temperatures are.

To avoid either extreme – and thus capture the actual population dynamics resulting from a given thermal regime – we calculated the average variance and deviation from the time-varying equilibrium of the population size for a given uncorrelated, normally-distributed thermal regime at time step lengths ranging from $\delta t$ = 0.01 to 100 (Fig. S2). Values of $\delta t$ where those mean quantities are changing the fastest – between where they level off at either extreme – are where accurate dynamics caused by the regime itself, and not the chosen time step size, are captured.

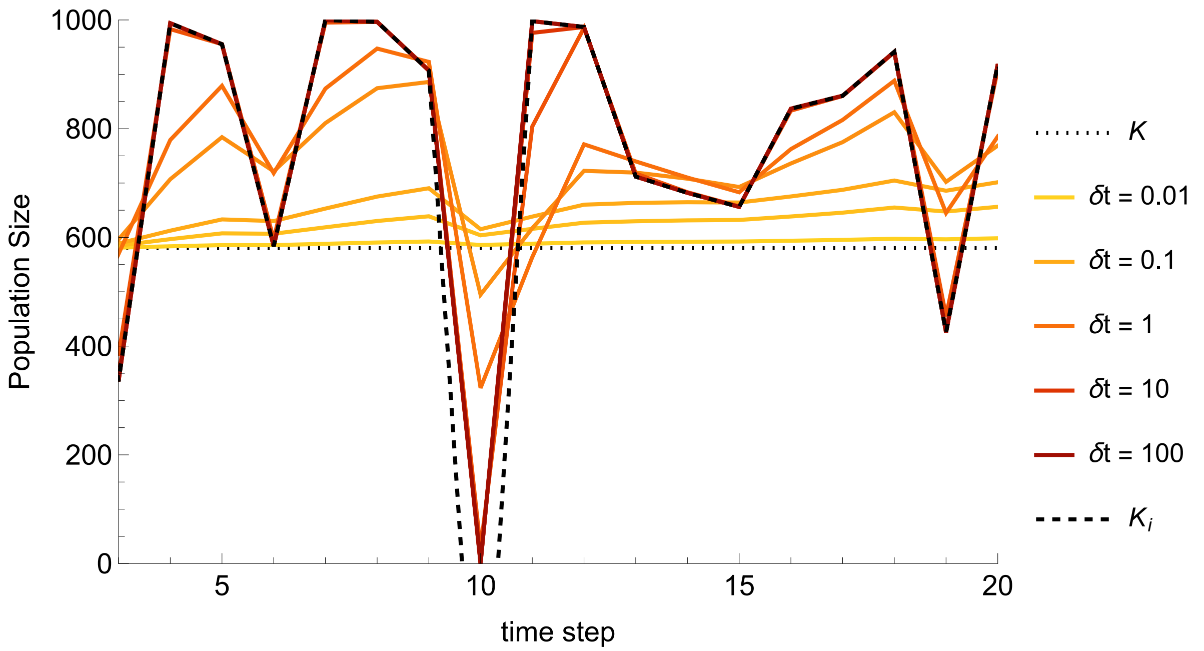

**Figure S1:** Sample plot of different population abundances calculated for the same sample thermal regime but with different time step lengths, compared to the constant equilibrium $K$ and the time-varying equilibrium $K_i$. The temperature regime is random and distributed according to $N(25, 3)$. In this example run, when $\delta t \geq 10$, the population goes extinct at time step 10.

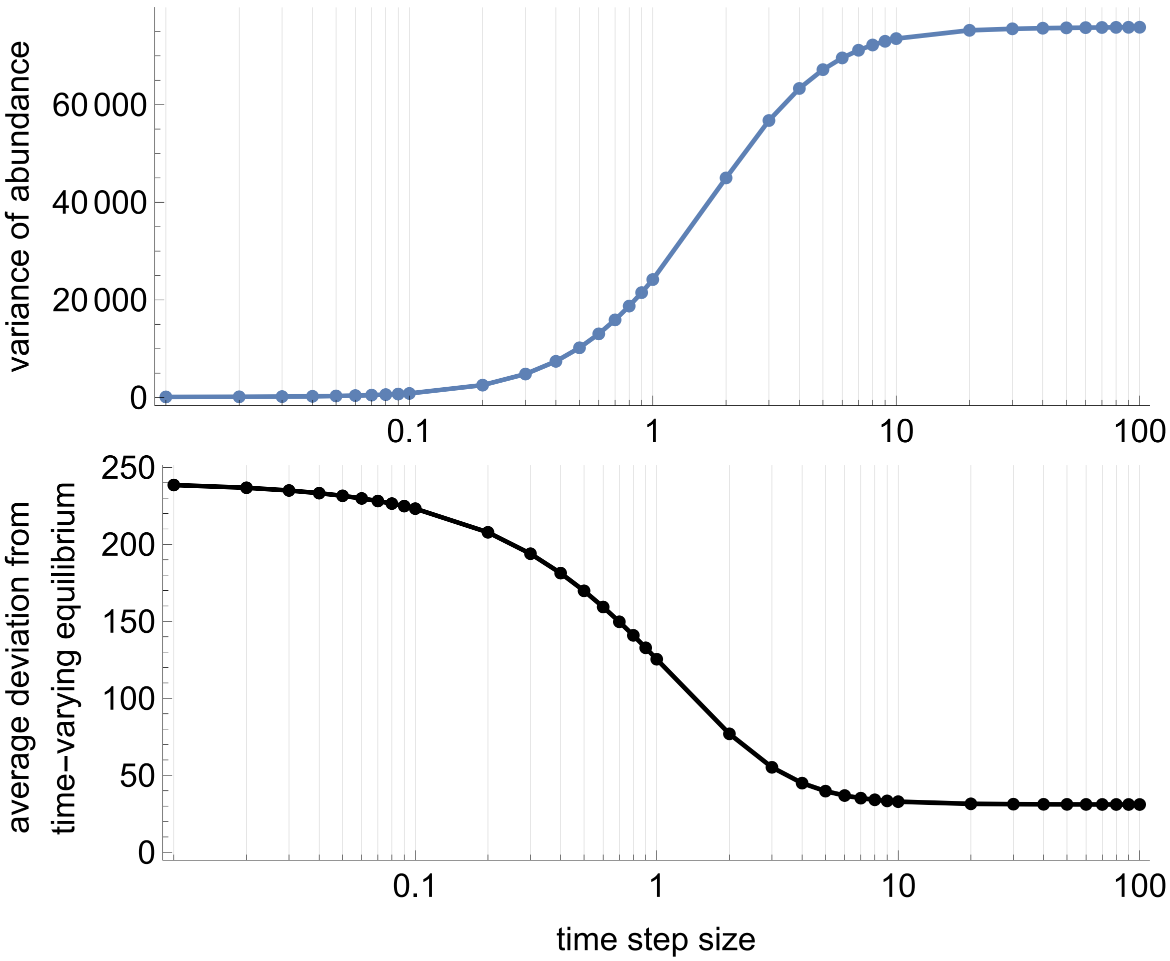

**Figure S2:** Given a time step size between 0.01 and 100, the average (a) variance of population abundance and (b) average deviation from the time-varying equilibrium Ki are calculated for a population undergoing the same thermal regime. At short time step sizes ($\delta t \leq 0.1$), slopes are near 0 and dynamics are dominated by the equilibrium state $K$ and the average fitness of the thermal regime. At long time step sizes ($\delta t \geq 10$), slopes are again near 0 and dynamics are dominated by the time-varying equilibrium Ki. Results show average values across 500 runs of 100 time steps each; the thermal regime of each run was unordered and randomly selected from $N(25,3)$.

**Appendix S5** *Extinction Risk Across the Thermal Landscape: Tropical Results*

The second model's results for the tropical, instead of the temperate, TPC. While there are quantitative differences in the proportion (Fig. S1) and timing (Fig. S2) of extinctions, the qualitative patterns are extremely similar. The primary quantitative difference for the tropical TPC is a slower ramping up of the proportion of extinctions as the thermal regime intensifies (e.g., no extinctions at $\sigma = 2$ and fewer between $2.5 < \sigma < 3.5$ in Fig. S1 (a)).

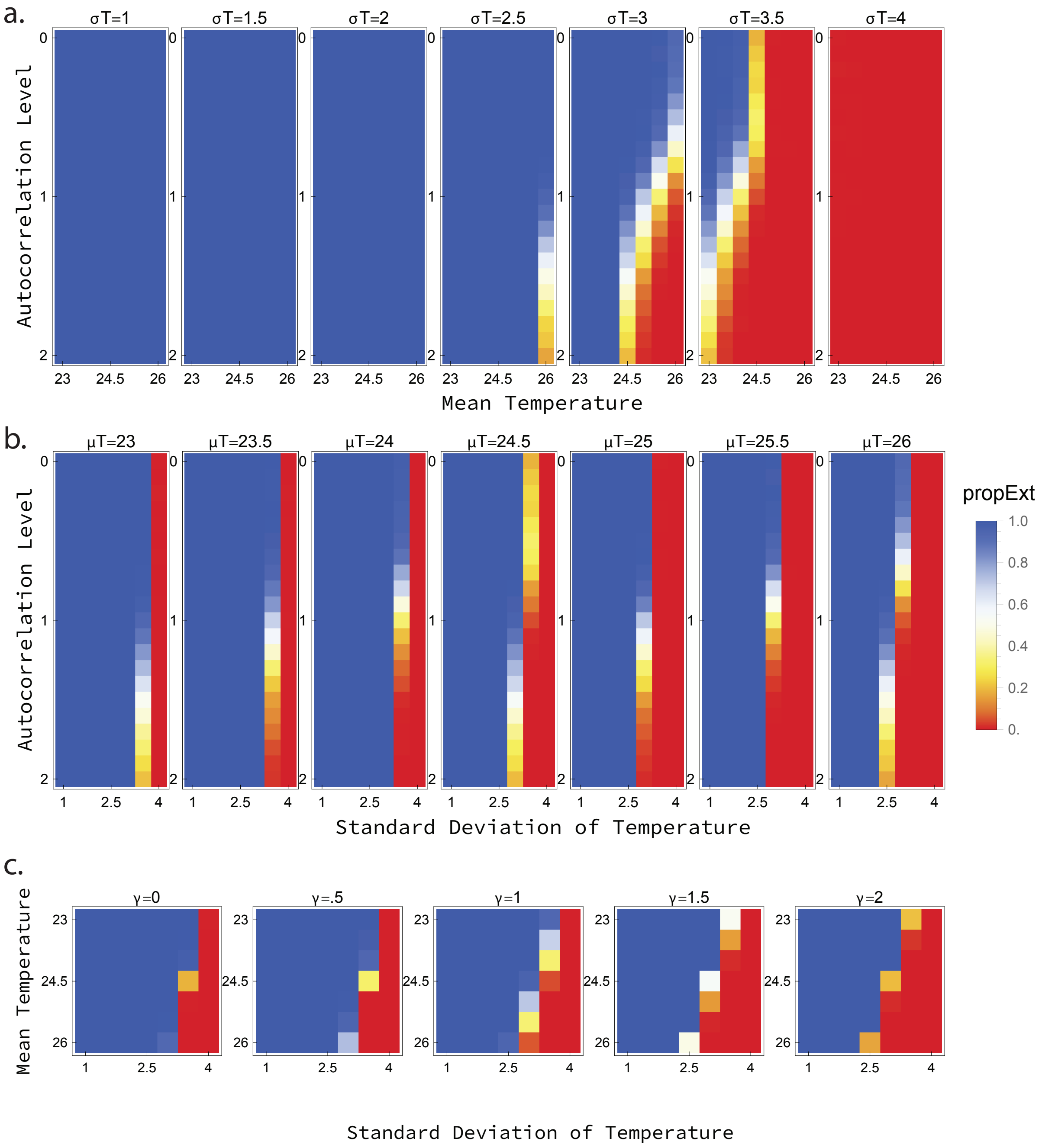

**Figure S1:** The proportion of runs going extinct at each autocorrelation level and thermal regime for the tropical TPC. (a) Mean temperature against autocorrelation level at each standard deviation. (b) Standard deviation against autocorrelation level at each mean temperature. (c) Standard deviation against mean temperature at each autocorrelation level.

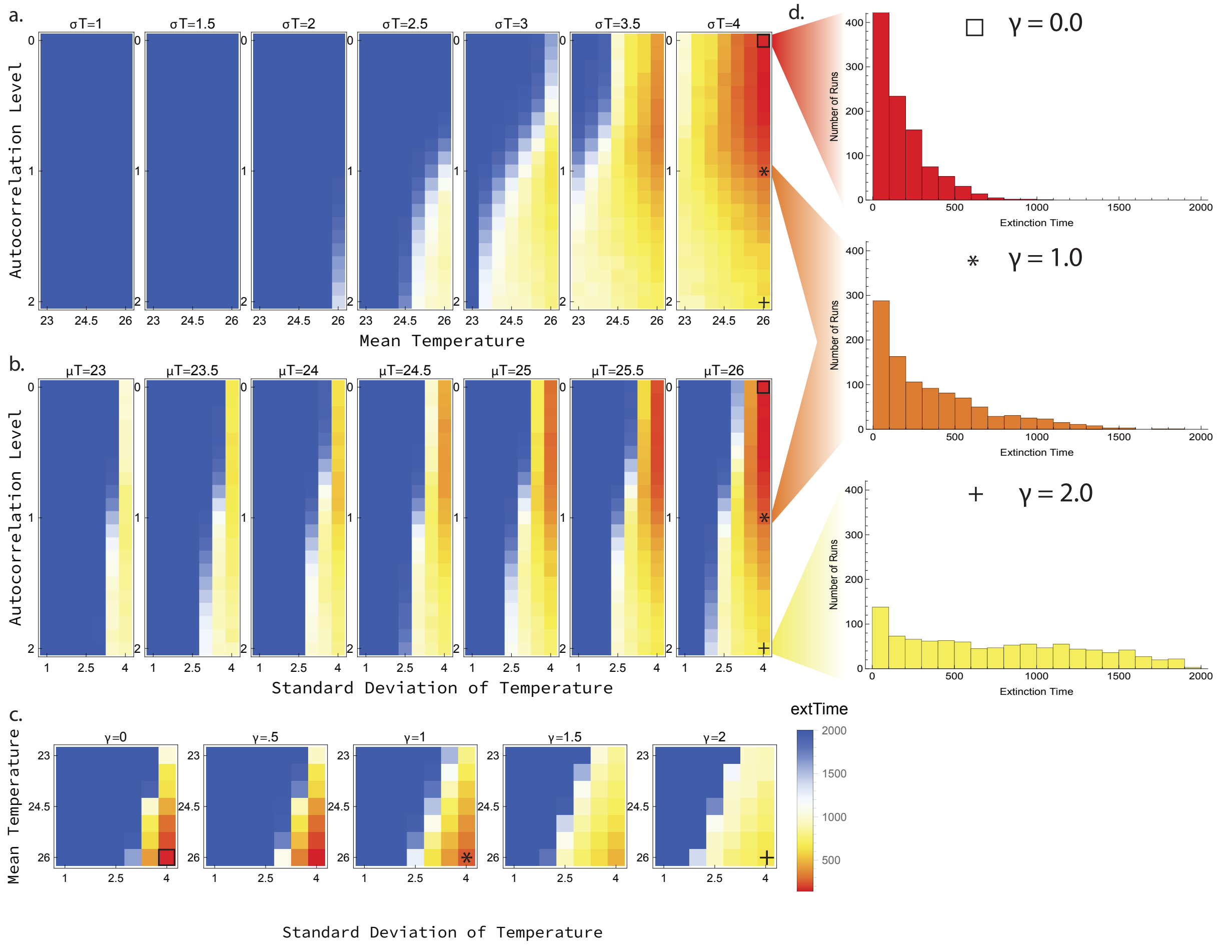

**Figure S2:** The average extinction time across runs for the tropical TPC at each autocorrelation level and thermal regime. (a) Mean temperature against autocorrelation level at each standard deviation. (b) Standard deviation against autocorrelation level at each mean temperature. (c) Standard deviation against mean temperature at each autocorrelation level. (d) Histograms of the extinction times at the highest mean and standard deviations of temperatures at different autocorrelation levels. Squares (γ = 0), stars (γ = 1), and plus signs (γ = 2) indicate the location of the histograms in each set of heat plots.

**Appendix S6** *Extended Model 3 Results*

**Figure S1:** The complete input and output data for each of the 38 species included in the final model. Note that axes are consistent across species for the left and right plots but vary based on distributions in the center plots.

*Left:* Gaussian-Quadratic TPC curve fit of the thermal maximum ($T_{\text{max}}$), thermal optimum ($T_{\text{opt}}$), and breadth parameter ($\sigma_p = \frac{T_{\text{opt}} - CT_{\text{min}}}{4}$). Graphs are colored by region according to the same scheme as Duffy et al. 2022, with purple indicating Northern Hemisphere Extra-tropics (30ºN to 90ºN), green indicating Tropics (30ºS to 30ºN), and orange indicating Southern Hemisphere Extra-tropics (90ºS to 30ºS).

*Center:* Histograms of the thermal data for each species' geographic coordinates (Visual Crossings Corporation, 2024), with historical temperature data (01-01-1994 to 12-31-2003) in yellow and recent temperature data (01-01-2014 to 12-31-2023) in blue. The mean temperatures (solid lines) plus and minus one standard deviation (dotted lines) are indicated in orange and blue, respectively.

*Right:* Plots of extinction risk for historic (gray) and recent (black) thermal distributions across autocorrelation levels with seasonal and diurnal fluctuations maintained across simulations. The actual spectral exponents of the observed temperatures – and the corresponding extinction risks – are indicated by the pink (historical) and red (recent) dots and dashed lines.

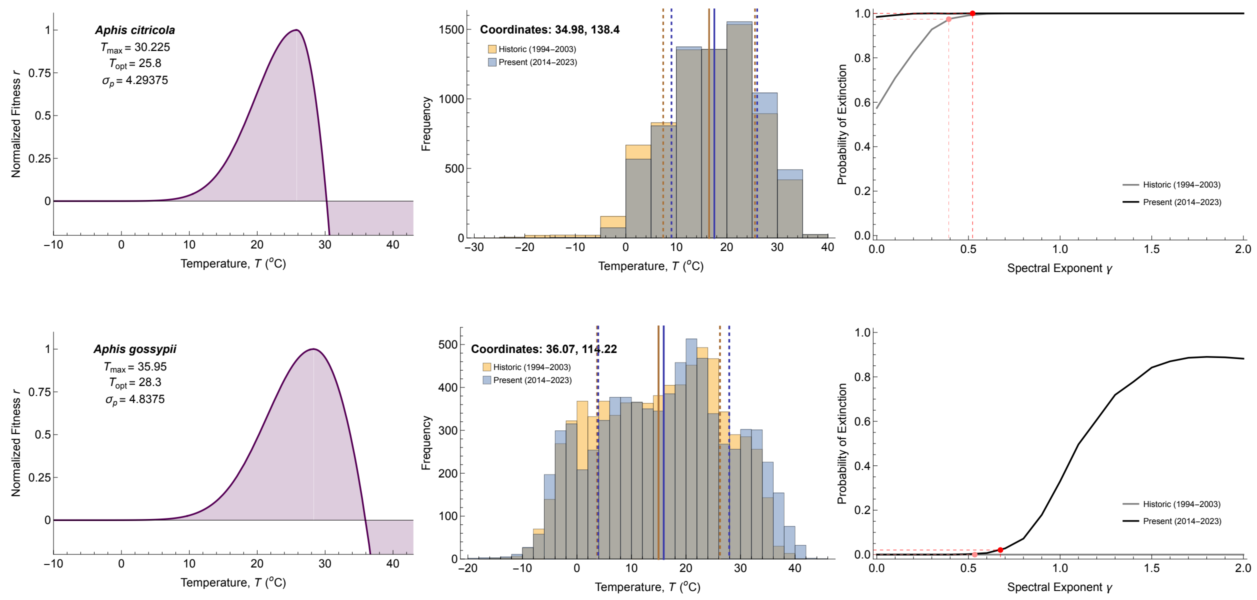

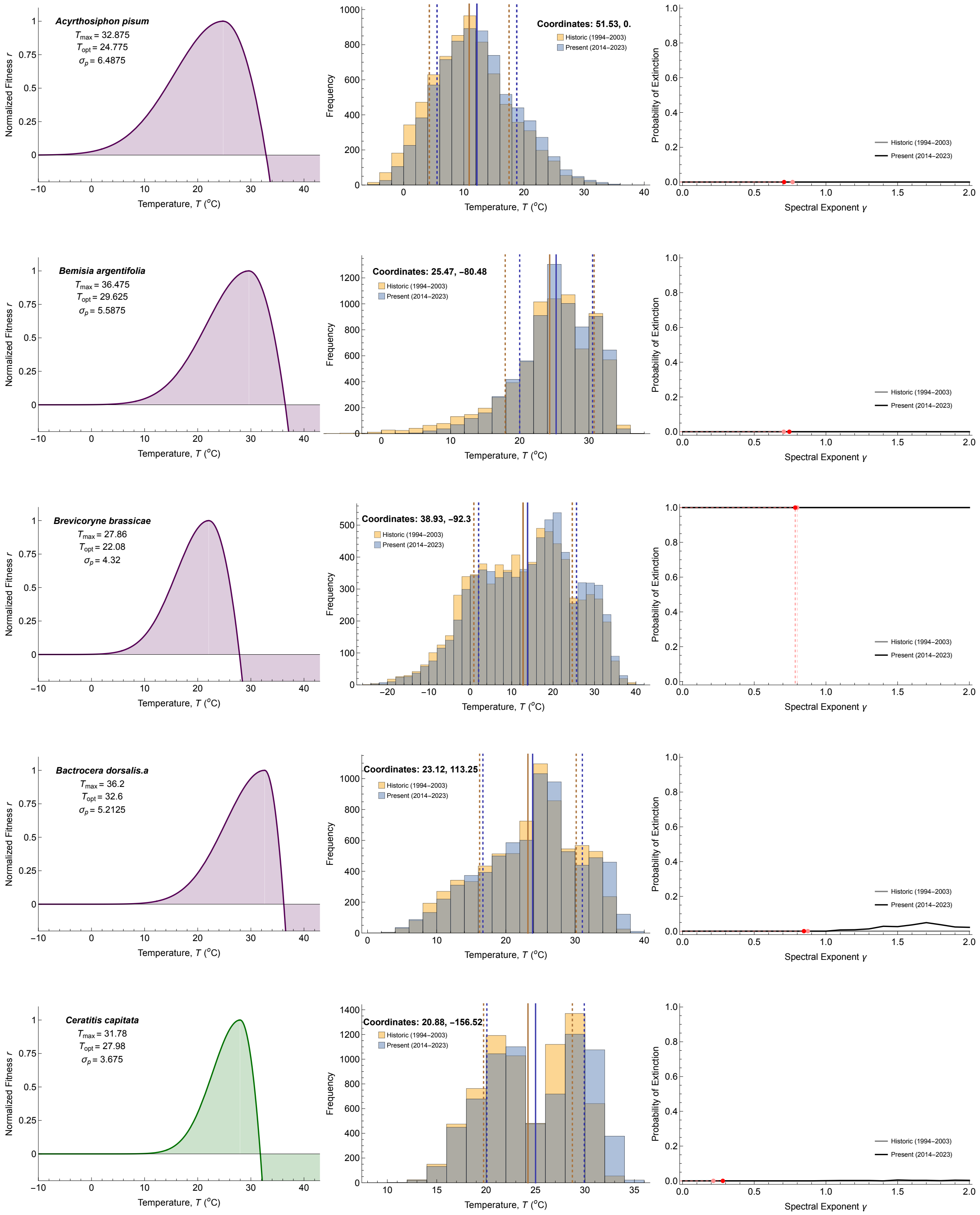

Acyrthosiphon pisum
T_max = 32.875
T_opt = 24.775
σ_p = 6.4875
Coordinates: 51.53, 0.
Bemisia argentifolia
T_max = 36.475
T_opt = 29.625
σ_p = 5.5875
Coordinates: 25.47, −80.48
Brevicoryne brassicae
T_max = 27.86
T_opt = 22.08
σ_p = 4.32
Coordinates: 38.93, −92.3
Bactrocera dorsalis.a
T_max = 36.2
T_opt = 32.6
σ_p = 5.2125
Coordinates: 23.12, 113.25
Ceratitis capitata
T_max = 31.78
T_opt = 27.98
σ_p = 3.675
Coordinates: 20.88, −156.52
Historic (1994–2003)
Present (2014–2023)
Normalized Fitness r
Temperature, T (°C)
Frequency
Probability of Extinction
Spectral Exponent γ

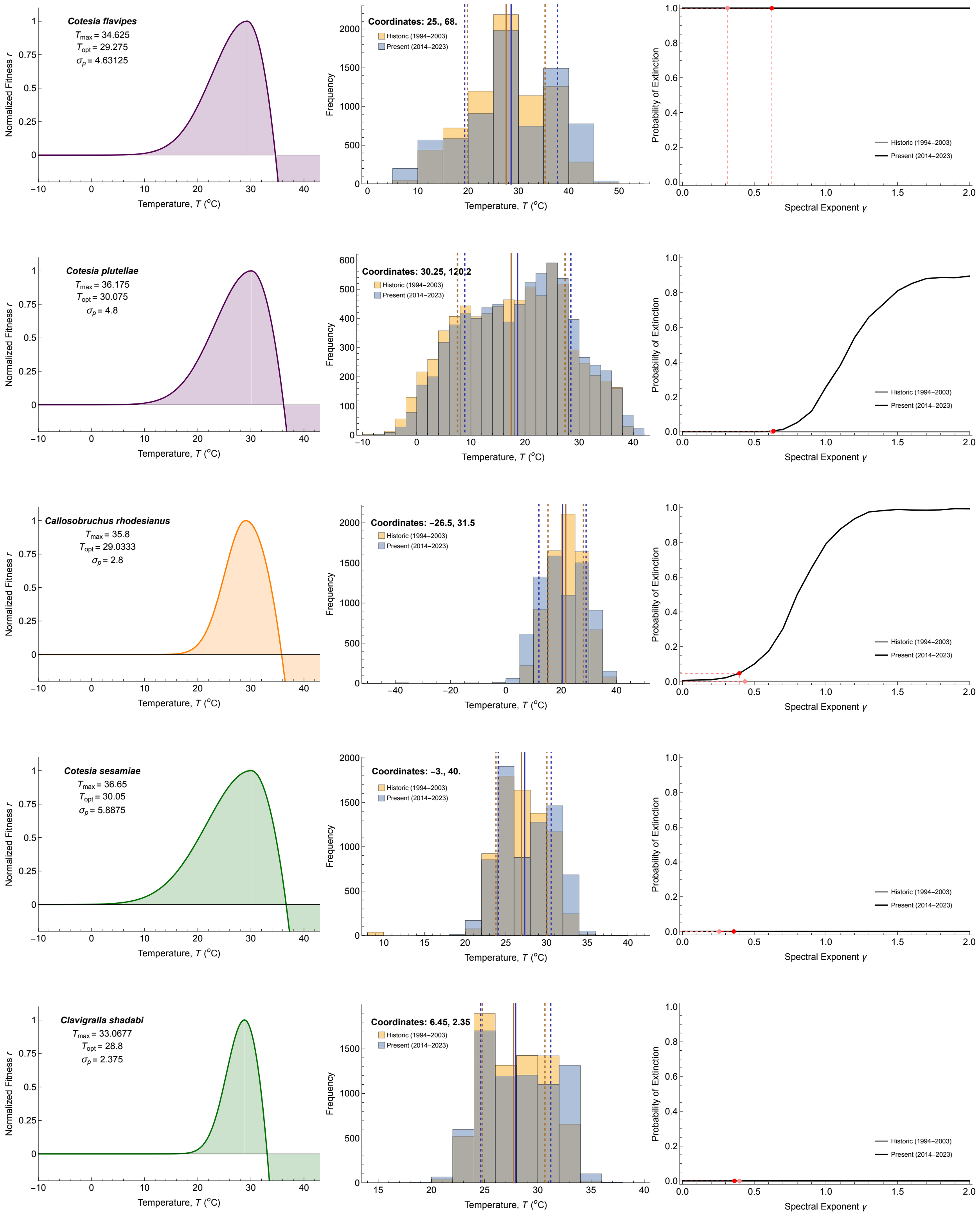
Cotesia flavipes
$T_{max}$ = 34.625
$T_{opt}$ = 29.275
$\sigma_p$ = 4.63125
Coordinates: 25., 68.
Cotesia plutellae
$T_{max}$ = 36.175
$T_{opt}$ = 30.075
$\sigma_p$ = 4.8
Coordinates: 30.25, 120.2
Callosobruchus rhodesianus
$T_{max}$ = 35.8
$T_{opt}$ = 29.0333
$\sigma_p$ = 2.8
Coordinates: −26.5, 31.5
Cotesia sesamiae
$T_{max}$ = 36.65
$T_{opt}$ = 30.05
$\sigma_p$ = 5.8875
Coordinates: −3., 40.
Clavigralla shadabi
$T_{max}$ = 33.0677
$T_{opt}$ = 28.8
$\sigma_p$ = 2.375
Coordinates: 6.45, 2.35
Historic (1994–2003)
Present (2014–2023)
Normalized Fitness r
Temperature, T (°C)
Frequency
Probability of Extinction
Spectral Exponent γ

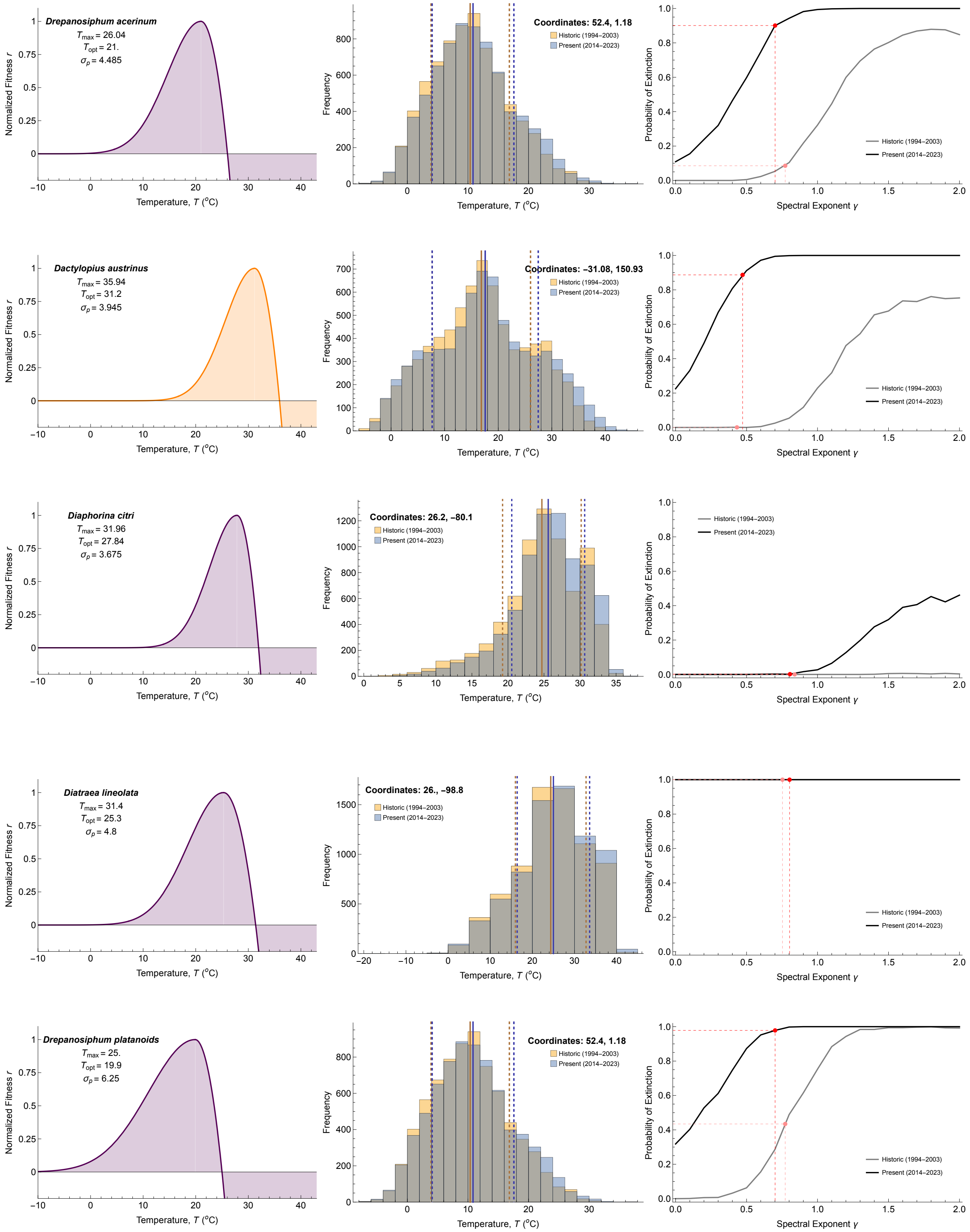
Drepanosiphum acerinum
$T_{max}$ = 26.04
$T_{opt}$ = 21.
$\sigma_p$ = 4.485
Coordinates: 52.4, 1.18
Historic (1994–2003)
Present (2014–2023)
Dactylopius austrinus
$T_{max}$ = 35.94
$T_{opt}$ = 31.2
$\sigma_p$ = 3.945
Coordinates: −31.08, 150.93
Diaphorina citri
$T_{max}$ = 31.96
$T_{opt}$ = 27.84
$\sigma_p$ = 3.675
Coordinates: 26.2, −80.1
Diatraea lineolata
$T_{max}$ = 31.4
$T_{opt}$ = 25.3
$\sigma_p$ = 4.8
Coordinates: 26., −98.8
Drepanosiphum platanoids
$T_{max}$ = 25.
$T_{opt}$ = 19.9
$\sigma_p$ = 6.25
Coordinates: 52.4, 1.18
Normalized Fitness r
Temperature, T (°C)
Frequency
Probability of Extinction
Spectral Exponent γ

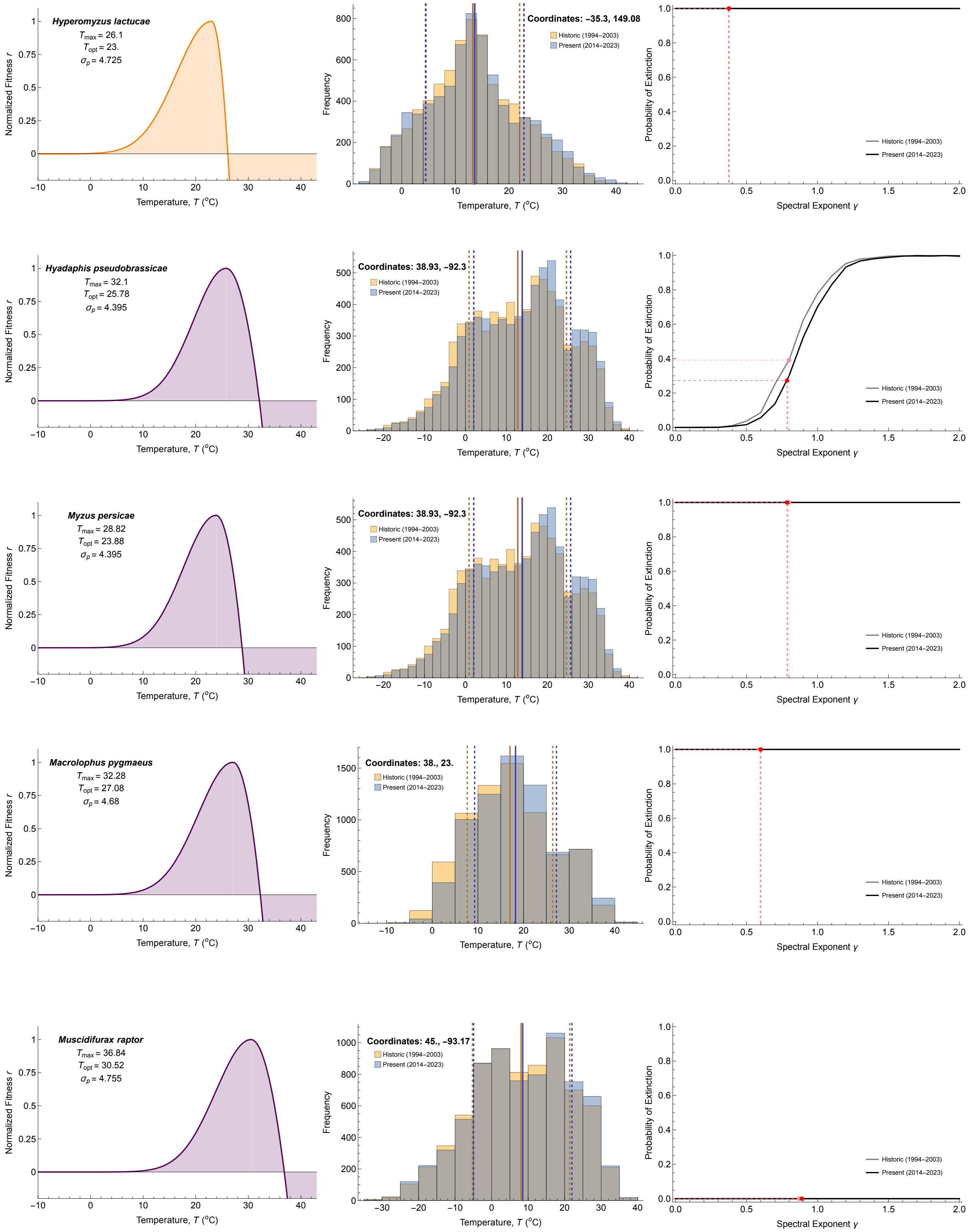
Hyperomyzus lactucae
$T_{max}$ = 26.1
$T_{opt}$ = 23.
$\sigma_p$ = 4.725
Coordinates: −35.3, 149.08
Hyadaphis pseudobrassicae
$T_{max}$ = 32.1
$T_{opt}$ = 25.78
$\sigma_p$ = 4.395
Coordinates: 38.93, −92.3
Myzus persicae
$T_{max}$ = 28.82
$T_{opt}$ = 23.88
$\sigma_p$ = 4.395
Coordinates: 38.93, −92.3
Macrolophus pygmaeus
$T_{max}$ = 32.28
$T_{opt}$ = 27.08
$\sigma_p$ = 4.68
Coordinates: 38., 23.
Muscidifurax raptor
$T_{max}$ = 36.84
$T_{opt}$ = 30.52
$\sigma_p$ = 4.755
Coordinates: 45., −93.17
Historic (1994–2003)
Present (2014–2023)
Normalized Fitness r
Temperature, T (°C)
Frequency
Probability of Extinction
Spectral Exponent γ

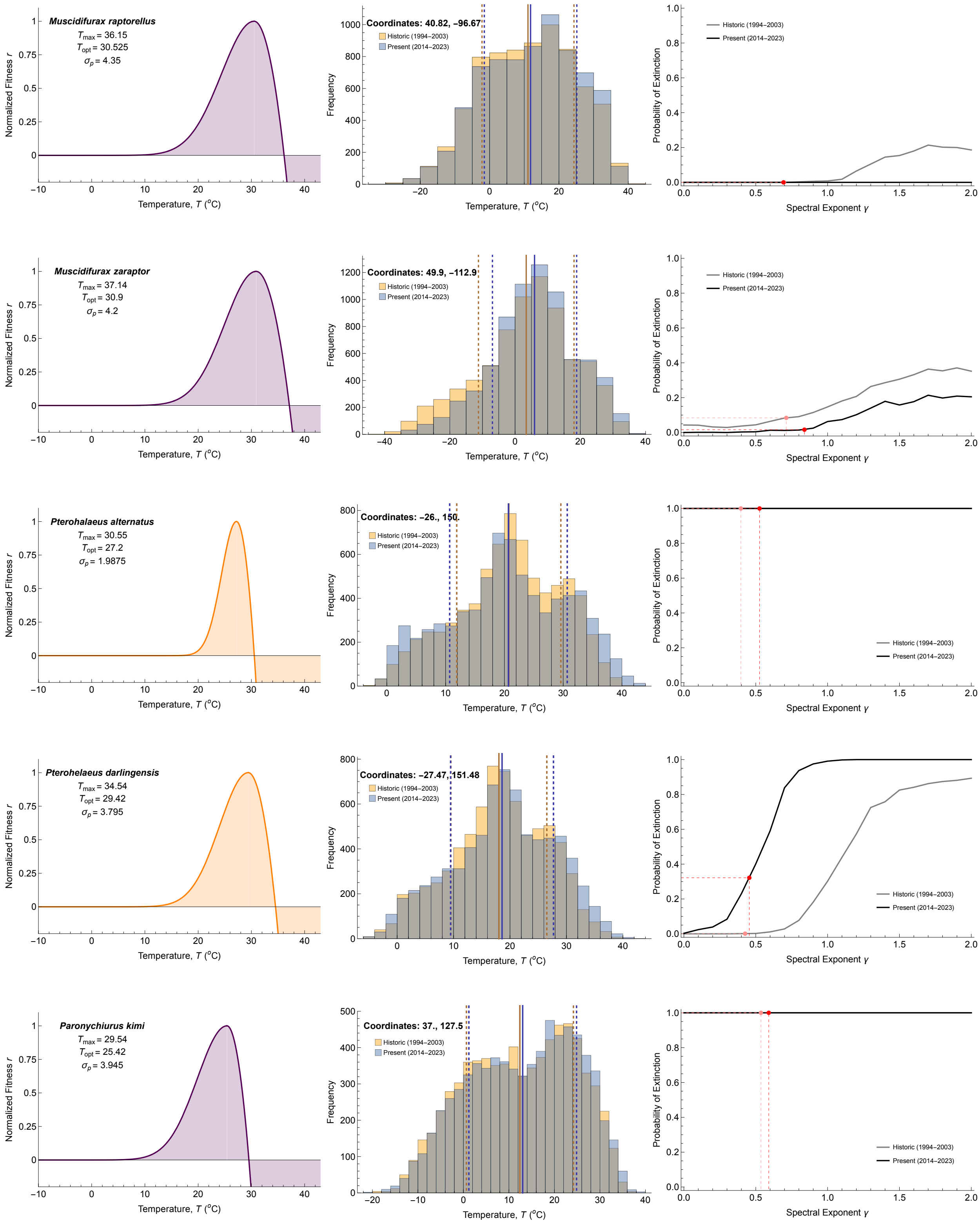
Muscidifurax raptorellus
$T_{max}$ = 36.15
$T_{opt}$ = 30.525
$\sigma_p$ = 4.35
Coordinates: 40.82, −96.67
Muscidifurax zaraptor
$T_{max}$ = 37.14
$T_{opt}$ = 30.9
$\sigma_p$ = 4.2
Coordinates: 49.9, −112.9
Pterohalaeus alternatus
$T_{max}$ = 30.55
$T_{opt}$ = 27.2
$\sigma_p$ = 1.9875
Coordinates: −26., 150.
Pterohelaeus darlingensis
$T_{max}$ = 34.54
$T_{opt}$ = 29.42
$\sigma_p$ = 3.795
Coordinates: −27.47, 151.48
Paronychiurus kimi
$T_{max}$ = 29.54
$T_{opt}$ = 25.42
$\sigma_p$ = 3.945
Coordinates: 37., 127.5
Historic (1994–2003)
Present (2014–2023)
Normalized Fitness r
Temperature, T (°C)
Frequency
Probability of Extinction
Spectral Exponent γ

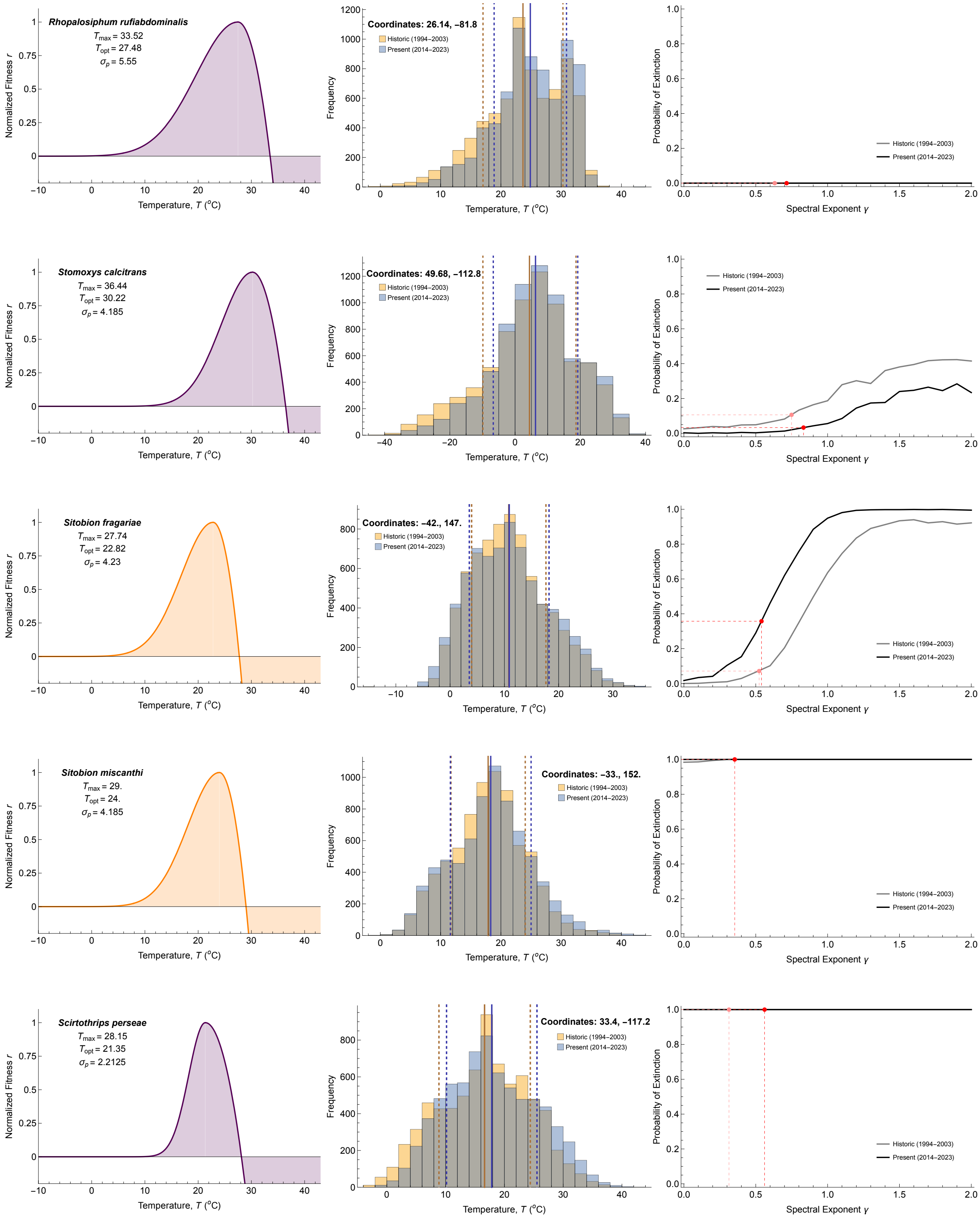

Rhopalosiphum rufiabdominalis
$T_{max}$ = 33.52
$T_{opt}$ = 27.48
$\sigma_p$ = 5.55
Coordinates: 26.14, −81.8
Stomoxys calcitrans
$T_{max}$ = 36.44
$T_{opt}$ = 30.22
$\sigma_p$ = 4.185
Coordinates: 49.68, −112.8
Sitobion fragariae
$T_{max}$ = 27.74
$T_{opt}$ = 22.82
$\sigma_p$ = 4.23
Coordinates: −42., 147.
Sitobion miscanthi
$T_{max}$ = 29.
$T_{opt}$ = 24.
$\sigma_p$ = 4.185
Coordinates: −33., 152.
Scirtothrips perseae
$T_{max}$ = 28.15
$T_{opt}$ = 21.35
$\sigma_p$ = 2.2125
Coordinates: 33.4, −117.2
Historic (1994–2003)
Present (2014–2023)
Normalized Fitness r
Temperature, T (°C)
Frequency
Probability of Extinction
Spectral Exponent γ

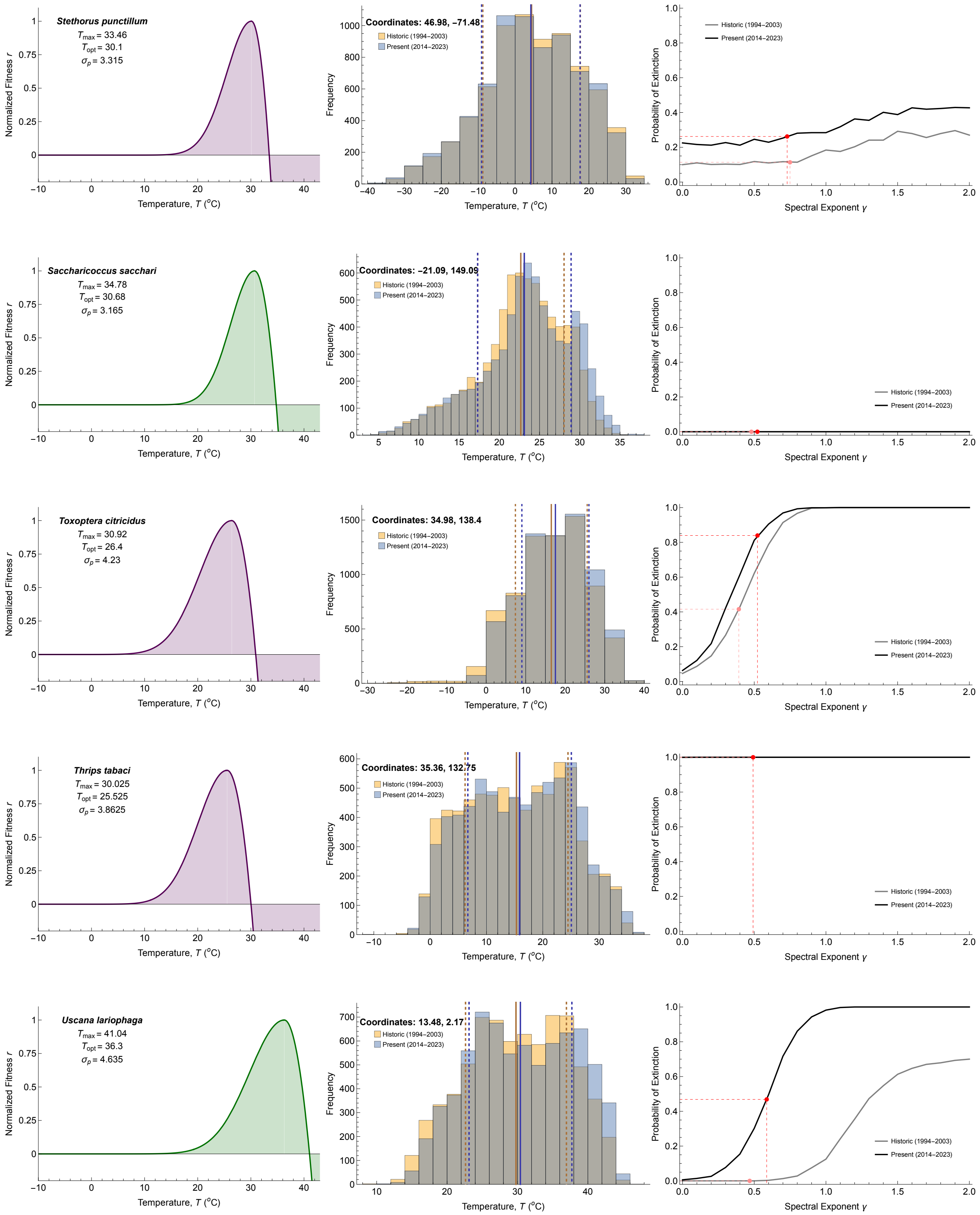

Stethorus punctillum
$T_{max}$ = 33.46
$T_{opt}$ = 30.1
$\sigma_p$ = 3.315
Normalized Fitness r
Temperature, T (°C)
Coordinates: 46.98, −71.48
Historic (1994–2003)
Present (2014–2023)
Frequency
Temperature, T (°C)
Probability of Extinction
Spectral Exponent γ
Historic (1994–2003)
Present (2014–2023)
Saccharicoccus sacchari
$T_{max}$ = 34.78
$T_{opt}$ = 30.68
$\sigma_p$ = 3.165
Coordinates: −21.09, 149.09
Toxoptera citricidus
$T_{max}$ = 30.92
$T_{opt}$ = 26.4
$\sigma_p$ = 4.23
Coordinates: 34.98, 138.4
Thrips tabaci
$T_{max}$ = 30.025
$T_{opt}$ = 25.525
$\sigma_p$ = 3.8625
Coordinates: 35.36, 132.75
Uscana lariophaga
$T_{max}$ = 41.04
$T_{opt}$ = 36.3
$\sigma_p$ = 4.635
Coordinates: 13.48, 2.17

**Appendix S7** *Risk assessments for populations with no change in probability of extinction*

For those species whose probability of extinction did not change between historical and present scenarios, we instead measured the mean time to extinction for species that always went extinct (Fig. S1) and mean minimum population size for species that never went extinct (Fig. S2).

**Figure S1:** Plots of the mean time to extinction across 1000 simulations at each autocorrelation level for species whose projected probability of extinction was 100% under both historical (gray) and recent (black) temperature regimes. The actual spectral exponents of the observed temperatures – and the corresponding extinction risks – are indicated by the pink (historical) and red (recent) dots and dashed lines. Included are three Southern Hemisphere species (*H. lactucae, S. miscanthi,* and *P.alternatus*), zero tropical species, and seven Northern Hemisphere species (*C. flavipes, D. lineolate, S. persea, T. tabaci, P. kimi, M.pygmaeus, M.persicae,* and *B. brassicae*).

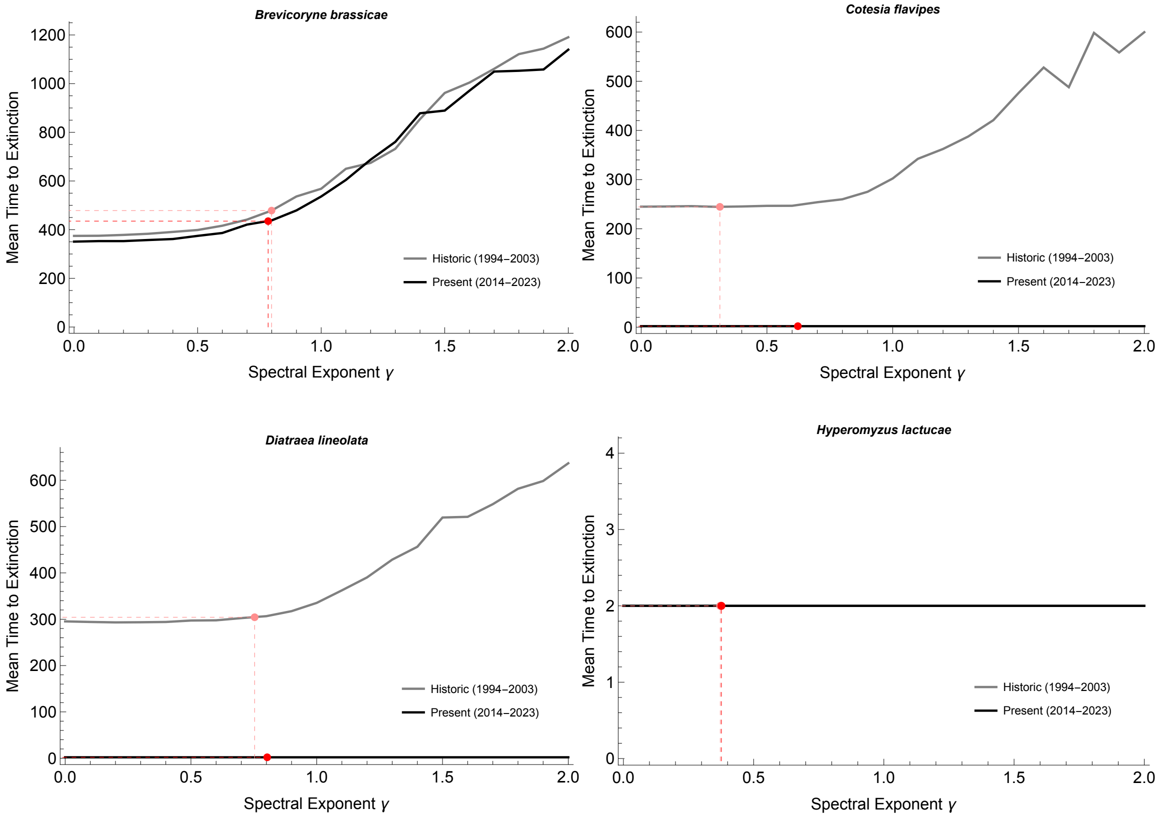

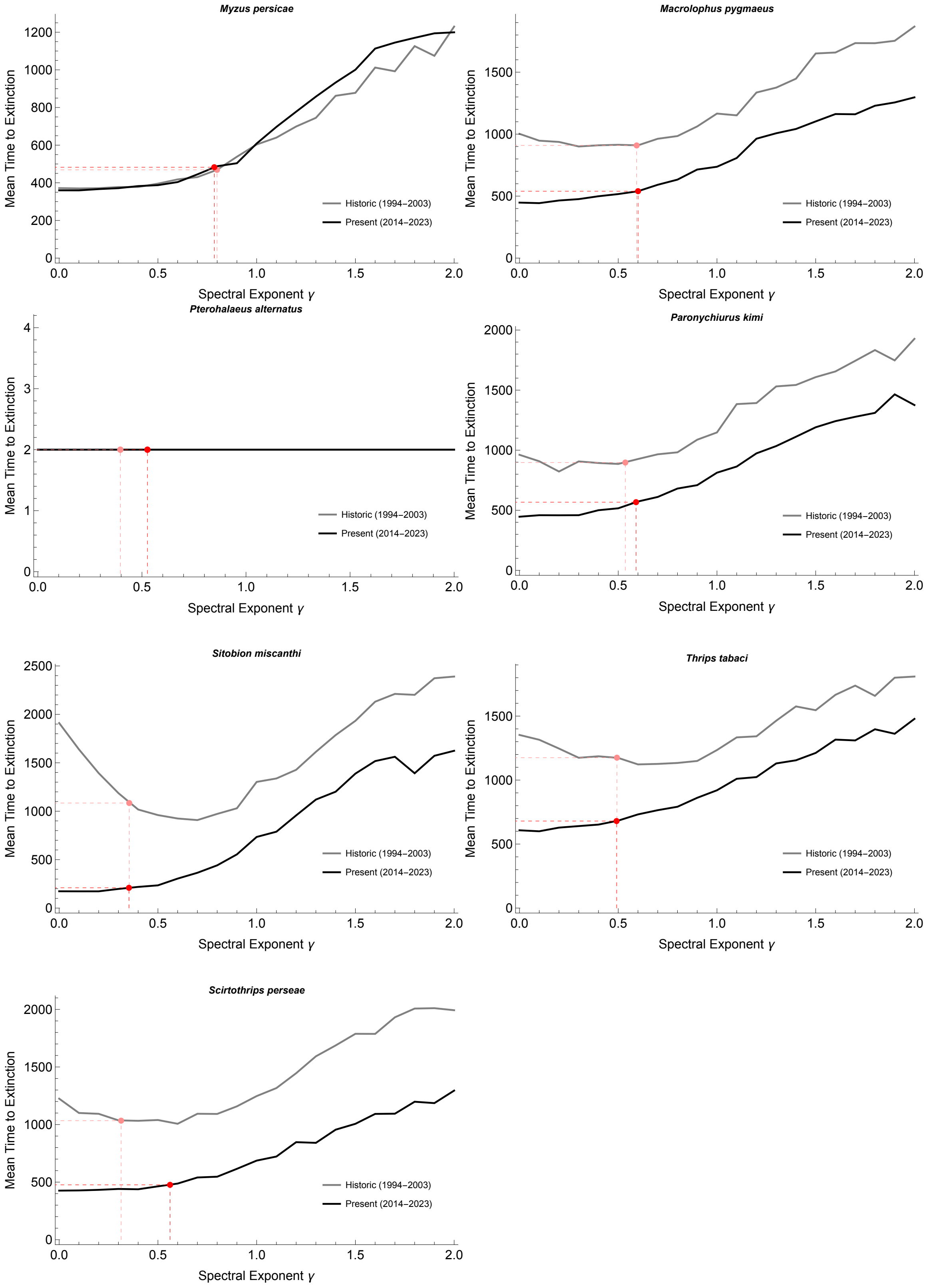
Myzus persicae
Macrolophus pygmaeus
Pterohalaeus alternatus
Paronychiurus kimi
Sitobion miscanthi
Thrips tabaci
Scirtothrips perseae
Mean Time to Extinction
Spectral Exponent γ
Historic (1994–2003)
Present (2014–2023)

**Figure S2:** Plots of average minimum abundance across 1000 simulations at each autocorrelation level for species whose projected probability of extinction was 0% under both historical (gray) and recent (black) temperature regimes. The actual spectral exponents of the observed temperatures – and the corresponding extinction risks – are indicated by the pink (historical) and red (recent) dots and dashed lines. Included are one Southern Hemisphere species (*E. lanigerum*), four Tropical species (*S. sacchari, C. sesamiae, C. shadabi,* and *C. capitata*), and six Northern Hemisphere species (*B. dorsalis, B. argentifolia, R. rufiabdominalis, M. rpatorellus, M.raptor,* and *A. pisum*).

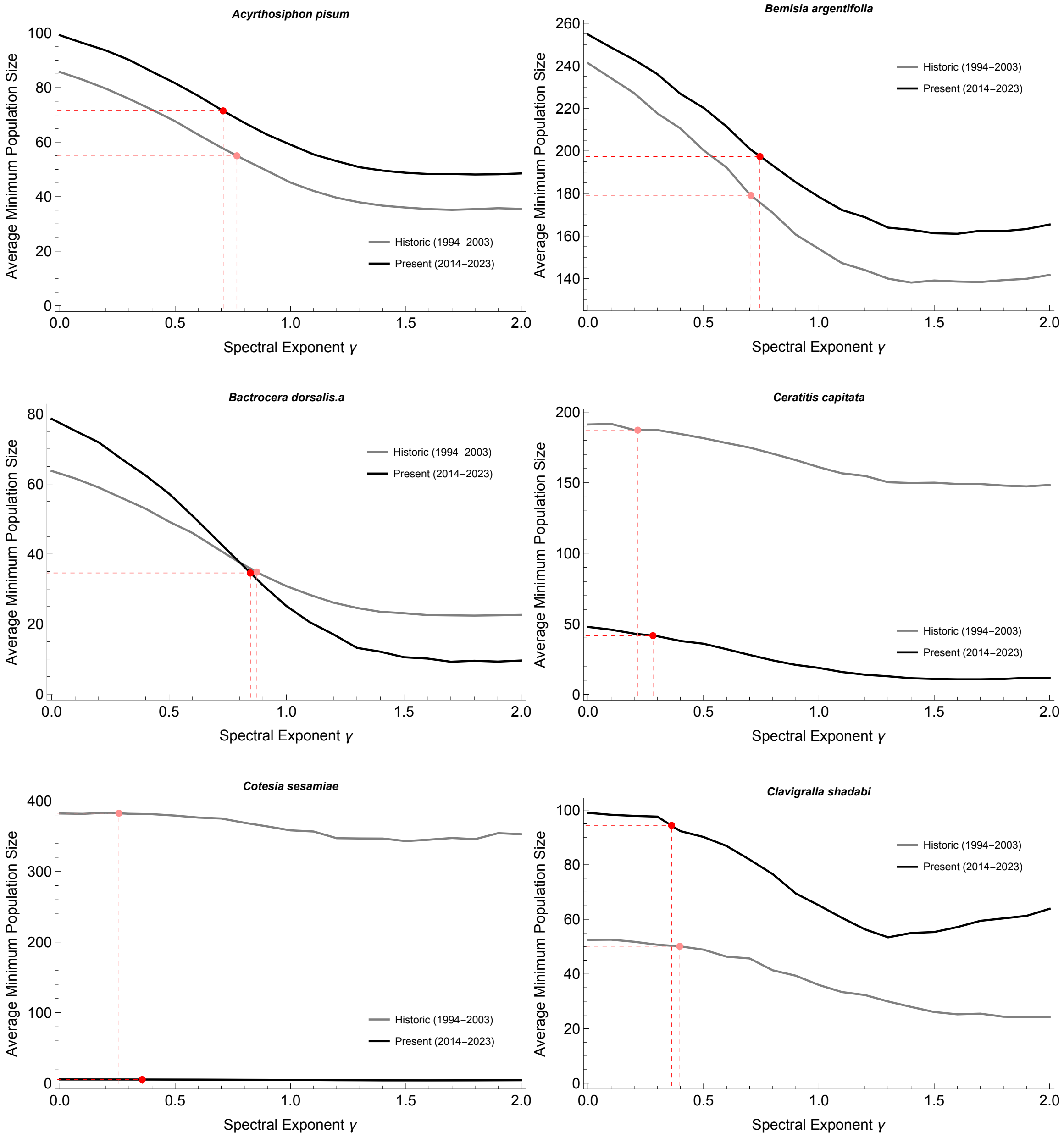

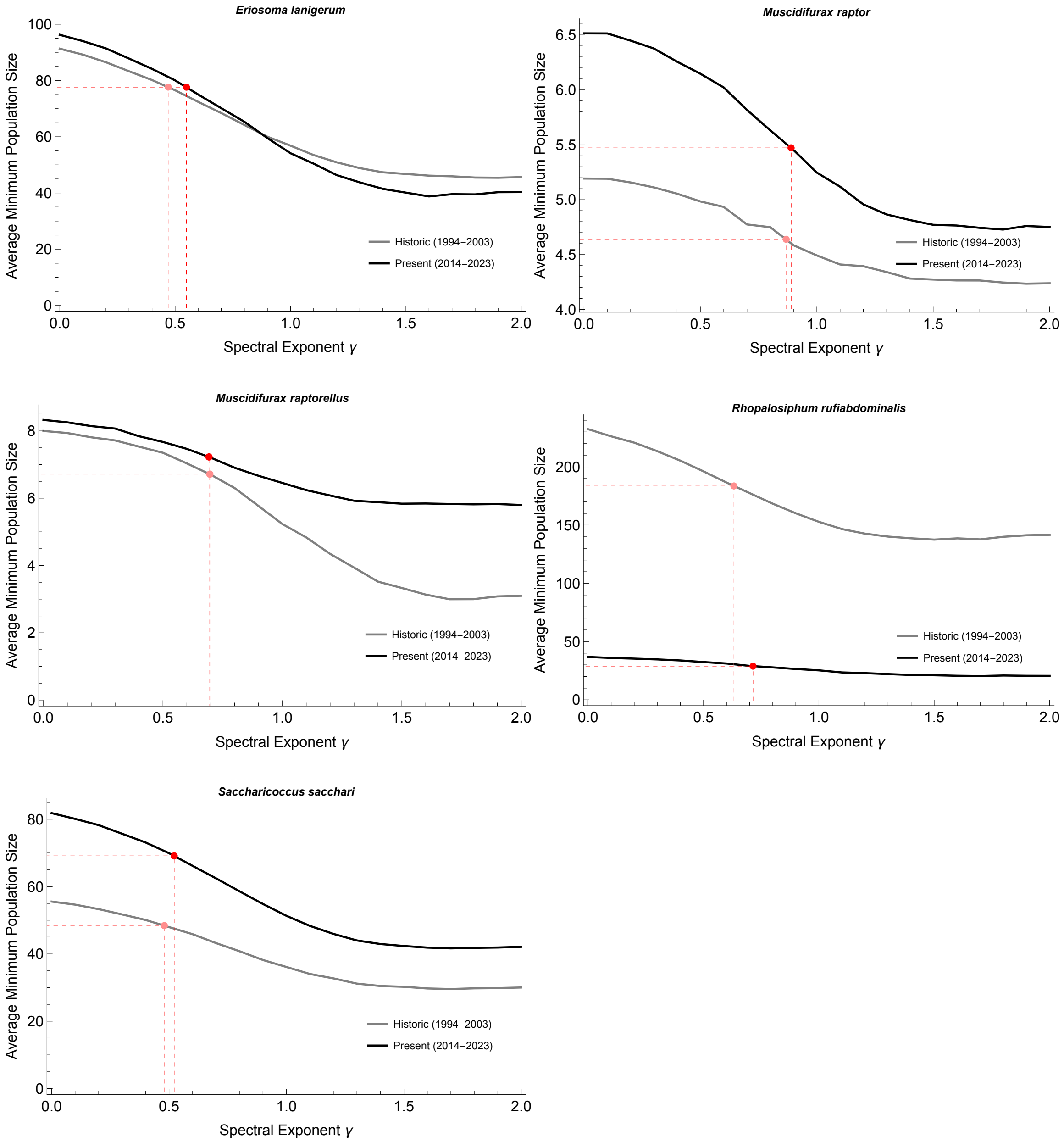
Eriosoma lanigerum
Muscidifurax raptor
Muscidifurax raptorellus
Rhopalosiphum rufiabdominalis
Saccharicoccus sacchari
Average Minimum Population Size
Spectral Exponent γ
Historic (1994–2003)
Present (2014–2023)
